# Evolution of the neural sex-determination system in insects: The *doublesex* homolog in crickets functions as an X-linked masculinizing factor to regulate morphological but not behavioral sexual dimorphism

**DOI:** 10.1101/2024.09.25.614005

**Authors:** Takayuki Watanabe

## Abstract

In animals, sex-specific social behaviors, such as courtship and aggression, are controlled by sex-specific neural circuits. To understand the neural mechanisms of sex-specific social behaviors from a comparative perspective, it is necessary to understand the sex determination mechanisms of neural circuits and their evolution across animal lineages. To address this, I have focused on the evolution of the neural sex determination system in insects. In the fruit fly *Drosophila melanogaster*, a model insect with powerful genetics, the molecular mechanisms underlying neuronal sex determination are well understood, and two transcription factor genes, *fruitless* and *doublesex* (*dsx*), have been identified as terminal differentiators. In contrast, those in direct-developing insects, which diverged from the holometabolous lineage, including *Drosophila*, about 400 million years ago, remain unclear. My previous studies found that the *fruitless* homolog is unlikely to be integrated into the neuronal sex determination system in direct-developing insects. In this study, I investigated whether the *dsx* homolog in direct-developing insects contributes to neuronal sex determination to form neural circuits that regulate adult social behaviors. I identified the *dsx* homolog in the two-spotted cricket *Gryllus bimaculatus* (*Gryllus dsx* gene) and revealed that it has a unique isoform configuration, and its encoded proteins contain a cryptic DNA-binding DM domain with accumulated amino acid substitutions. A knockout study revealed that the *Gryllus dsx* gene functions as an X-linked morphological masculinizing factor but does not determine the sexual orientation or behavioral patterns of adult social behavior in crickets. These results suggest that, at least in crickets, an unknown sex determination factor(s) other than *fruitless* and *dsx* plays a central role in neuronal sex determination.

## Introduction

In animals, sex-specific social behaviors, such as mating, courtship, and aggression, are regulated by sex-specific neural circuits. While both sexes share a common neural circuit configuration, these sex-specific circuits modulate brain function to produce behaviors characteristic of each sex. Consequently, identifying sex-specific neural circuits is crucial for understanding the neuronal mechanisms that underlie sex-specific brain functions (Roggenbuck et al., 2024). As the most diverse animal taxon with a history of over 500 million years of evolution, insects display a range of sex-specific social behaviors, including varied courtship rituals, maternal care, and male aggression over females or territories. Moreover, insects have exhibited significant flexibility in their postembryonic development, transitioning from ametabolous to holometabolous metamorphosis throughout their evolutionary history. These distinctive features make insects an invaluable model for investigating the neural basis of sex-specific social behavior, as well as the developmental and evolutionary processes that shape these behaviors.

Sex determination in insects is cell-autonomous (Bear & Monteiro, 2013), with each cell determining its own sex through a sex determination cascade (Baker, 1989; Yamamoto et al., 1998; Shirangi & McKeown, 2007). In *Drosophila*, the splicing factor gene *Sex lethal* (*Sxl*), which acts as the master switch for female development, is transcribed under conditions where the ratio of the number of X chromosomes to autosomes is greater than 1 (Cline, 1993). Sxl then regulates the splicing of its downstream target *transformer* (*tra*) to produce functional Tra protein in female cells (Inoue et al., 1990). The female-specific functional Tra protein (Tra^F^) functions as an RNA-binding protein, forming a female-specific splicing factor complex with Transformer-2 (Tra-2) and RBP1 proteins (Hoshijima et al., 1991; Heinrichs & Baker, 1995). The Tra^F^/Tra-2 complex then regulates the splicing of two transcription factor genes, *fruitless* and *doublesex* (*dsx*), the terminal differentiators in the *Drosophila* sex determination cascade (Coschigano & Wensink, 1993; Drapeau et al., 2003), to switch gene expression in cells to the female form. In male cells where Sxl is not expressed at the initial step of the cascade, the *tra* gene is spliced to produce a male-specific non-functional truncated Tra protein (Tra^M^), which is not complex with Tra-2 and RBP1. As a result, the splicing of the *fruitless* and *dsx* is regulated in a male-specific manner (Yamamoto et al., 1998). In the adult *Drosophila* brain, neurons expressing sex-specific gene products of *fruitless* and *doublesex* form sexually dimorphic circuits, which regulate sexually dimorphic behaviors including courtship behavior and aggression (Manoli et al., 2005; Demir & Dickson, 2005; Vrontou et al., 2006; Chan & Kravitz, 2007; Rideout et al., 2007; Kimura et al., 2008; Rideout et al., 2010; Dauwalder, 2011; Shirangi et al., 2016; Ishii et al., 2020; Han et al., 2022; Sato & Yamamoto, 2023).

Although morphological sex differences in the brain have been reported in various insect species (e.g., Rospars & Hildebrand, 2000; Nishikawa et al., 2008; Jundi el et al., 2009; Nishino et al., 2009; Hu et al., 2011), the morphological and functional sex differences in individual neural circuits are hardly examined, except in *Drosophila*, where powerful genetic tools are available (Baker et al., 2024). Researchers, therefore, infer the molecular mechanisms of neuronal sex determination in non-Drosophila insects from gene structure and gene expression, as well as from analysis of the behavioral phenotype when gene function is suppressed. For example, *fruitless* transcripts encoding the Fru^M^ homolog were found in several dipteran insects, which were produced under the control of the sex-specific splicing cascade (Gailey et al., 2006; Meier et al., 2013; Salvemini et al., 2013). In two hymenopteran insects (*Nasonia vitripennis* and *Apis mellifera*), sex-specific *fruitless* transcripts are produced by sex-specific alternative splicing (Bertossa et al., 2009). Interestingly, in Hymenoptera, the female-specific transcript encodes functional Fruitless protein (Fru^F^) with the BTB and zinc finger domains (Bertossa et al., 2009). The functional Fruitless isoforms (i.e., dipteran Fru^M^ and hymenopteran Fru^F^ proteins) contain an N-terminal extension adjacent to the BTB domain, whereas the non-functional Fruitless isoform of the other sex terminates translation after the N-terminal extension sequence. Besides, I have conducted a series of studies in the two-spotted cricket *Gryllus bimaculatus*, a direct-developing insect classically used in neuroethological studies, to elucidate the molecular and cellular mechanisms of neuronal sex determination in basal direct-developing insects.

Previously, I examined whether the *Gryllus fruitless* homolog was involved in neuronal sex determination in crickets (Watanabe, 2019). The *Gryllus fruitless* homolog does not have a P1-like promoter far distal to the gene body, nor does it possess an isoform generation mechanism dependent on alternative promoter usage and alternative splicing. This suggests that the *Gryllus fruitless* homolog lacks the mechanism for expressing sex-specific protein isoforms dependent on the P1 promoter, which is found in *Drosophila* and Hymenoptera (Ryner et al., 1996; Bertossa et al., 2009). Additionally, based on interspecies comparisons of gene structure, it was expected that the *fruitless* homologs of direct-developing insects would have a gene organization similar to that of the *Gryllus fruitless* homolog. With the above findings, I proposed an evolutionary model of the insect *fruitless* homolog: *fruitless* homologs did not originally function in the sex determination mechanism. However, in the common ancestor of holometabolous insects, *fruitless* homologs acquired the P1 promoter, and the P1 transcript acquired a target motif for sex-specific splicing. These two evolutionary events incorporated the *fruitless* homolog into the sex determination system as an additional terminal differentiator (Watanabe, 2019). This hypothesis leads us to expect that the prototypic neuronal sex determination system in insects would be dependent on *doublesex*, another terminal differentiator in the neuronal sex determination system in *Drosophila*.

The *Drosophila dsx* gene was first identified through a recessive mutation that transforms both males and females into an intersex phenotype (Hildreth, 1965). Subsequently, the genomic region of the *dsx* locus was cloned, revealing that the *dsx* transcript undergoes sex-specific splicing mediated by Tra/Tra2 (Baker & Wolfner, 1988; Burtis & Baker, 1989). The Dsx proteins were then characterized as zinc finger-type DNA-binding proteins (Erdman & Burtis, 1993). In parallel with studies on *Drosophila*, studies on sex determination systems in other animal lineages highlighted the presence of a conserved sex determination factor known as the DMRT (<u>D</u>oublesex and <u>M</u>ab-3-related transcription factor) family across Metazoa (Murphy et al., 2015; Mawaribuchi et al., 2019), and *Drosophila dsx* was the first member of this gene family. In addition to the prominent morphological phenotype of *dsx* mutants, early behavioral studies failed to detect behavioral abnormalities in *dsx* mutants (McRobert & Tompkins, 1985), leading to the conclusion that *dsx* was primarily a somatic sex determination factor. However, subsequent functional analysis of *dsx*-expressing neurons revealed that *dsx*-expressing neurons control sexual orientation, courtship song, male copulation, and aggression in *Drosophila* (Shirangi et al., 2016; Pavlou et al., 2016; Koganezawa et al., 2016; Peng et al., 2022; Han et al., 2022). Furthermore, studies on *dsx* homologs in non-*Drosophila* insects have revealed that the *dsx* gene regulates morphological sex determination, as well as sex-related differences in behavior in various insect species (Hu et al., 2011; Ito et al., 2013; Kunte et al., 2014; Gotoh et al., 2016; Iijima et al., 2019; Wexler et al., 2019; Singh Brar et al., 2022; Chikami et al., 2022). In this study, I conducted a series of molecular biological, biochemical, and genetic studies to investigate the contribution of the *Gryllus dsx* homolog to sex-specific behavior in crickets and the underlying sexual dimorphism in the nervous system.

## Materials and Methods

### 1. Animals

Two-spotted cricket *Gryllus bimaculatus* (De Geer) was reared in groups on a 14-hour light/10-hour dark cycle at 28°C. They were fed a diet of insect food pellets or basic food for hamsters (MF) (Oriental Yeast Co., Tokyo, Japan) and provided with water *ad libitum*. A wild-type strain (Hokudai WT strain; Watanabe et al., 2018) was used for experiments other than CRISPR-Cas9-based genome editing. A white-eye strain (Hokudai *gwhite* strain; Watanabe et al., 2018) was used for CRISPR-Cas9-based genome editing. All experiments were conducted during the light phase of the animals’ light cycle.

Camel cricket *Diestrammena* sp., tree cricket *Oecanthus longicauda,* and mayfly *Ephemera strigata* were collected in Sapporo (Hokkaido, Japan), while green tree cricket *Truljalia hibinonis* was collected in Hayama (Kanagawa, Japan). House cricket *Achita domestica*, katydid *Gampsocleis buergeri*, mole cricket *Gryllotalpa orientalis*, migratory locust *Locusta migratoria*, bell cricket *Meloimorpha japonica*, scaly cricket *Ornebius bimaculatus*, black field cricket *Teleogryllus commodus*, and *Vescelia pieli ryukyuensis* were purchased from local pet shops. The adult fruit fly *Drosophila melanogaster* (Canton S strain) was kindly provided by Dr. Nobuaki Tanaka (Hokkaido University).

### 2. Identification of *Gryllus dsx* gene expressed in the cricket brain

#### Homology search

To identify *dsx* homologs in crickets, a tBLASTn search was performed on the transcriptome shotgun assembly (TSA) of *Teleogryllus commodus* (BioProject ID: PRJNA252786) using Dsx protein sequences from various insect species as queries. As a result, an assembled transcript encoding a protein resembling other insect Dsx protein were found (GenBank ID: GBHB01043004).

#### RT-PCR-based cloning

To confirm the presence of a *dsx* homolog in the brain of *Gryllus bimaculatus*, a partial cDNA encoding the DM domain was amplified using primers designed based on the nucleotide sequence of the *Teleogryllus commodus* TSA. RT-PCR was performed according to the experimental procedure described in Watanabe et al. (2018). Subsequently, 5 ‘and 3 ‘rapid amplification of cDNA ends (RACE) was conducted to obtain the 5 ‘and 3 ‘regions of the gene. The full-length open reading frame (ORF) of the gene was then amplified using primers designed for the 5 ‘and 3 ‘untranslated regions (UTRs). The 5 ‘and 3 ‘RACE were performed using the FirstChoice RLM-RACE kit (Ambion, Austin, TX, USA), according to Watanabe et al. (2018). All PCRs were performed using the Q5 High-Fidelity DNA polymerase (New England Biolabs, MA, USA). Amplified cDNA fragments were cloned into the pGEM-T Easy vector (Promega, WI, USA). The nucleotide sequences of the obtained cDNAs were registered in GenBank (GenBank IDs: *Gryllus dsx^COM^*, LC841920; *Gryllus dsx^M^*, LC841922; *Gryllus dsx^F^*, LC841921). Primers used for 5 ‘and 3 ‘ RACEs and full-length cDNA amplification are listed in Table 1-1.

**Table 1-1.**
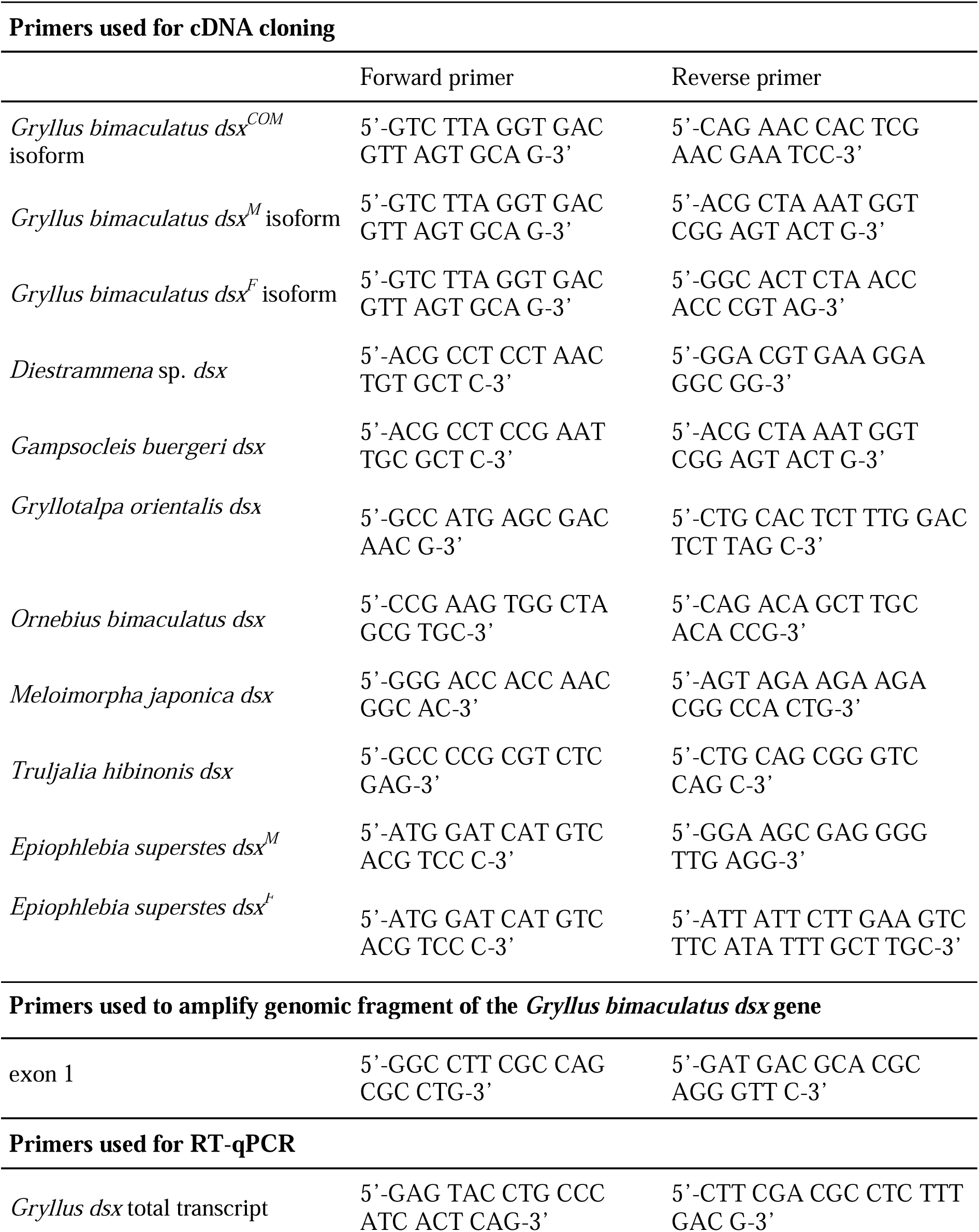
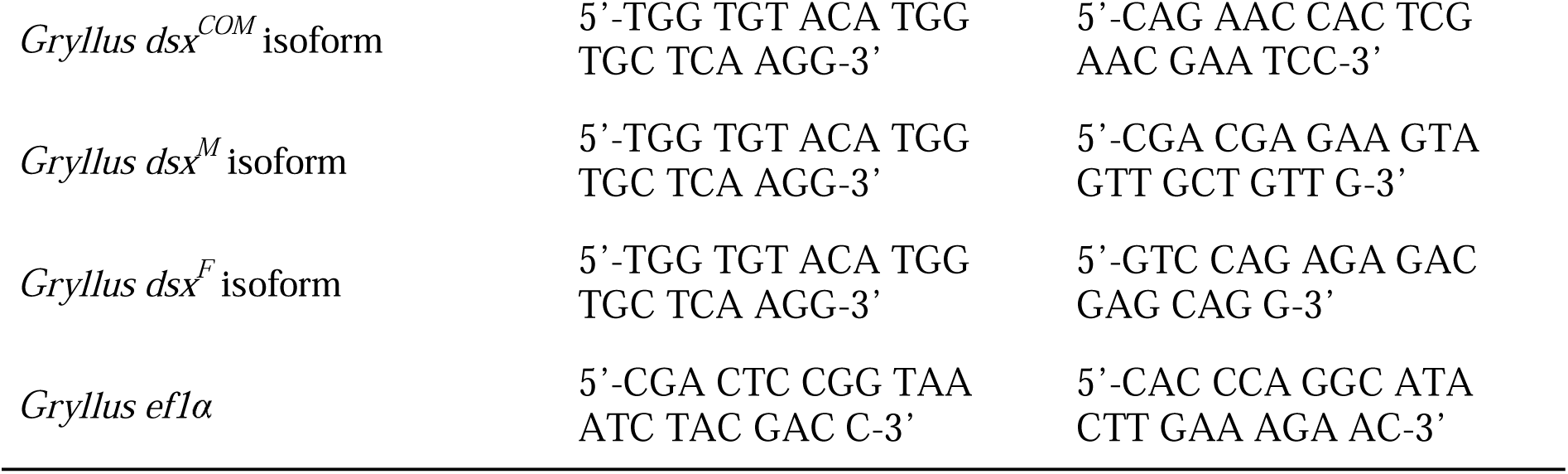
List of primers used for cloning and RT-qPCR.

### 3. Determination of the gene structure of the *Gryllus dsx* gene

Since the coding exon(s) of the 5 ‘region of the *Gryllus dsx* gene are missing in the current cricket genome assembly (Gbim_1.0; BioProject ID: PRJDB10609), inverse PCR was performed to obtain the genomic flanking regions of the missing exon(s) and determine the genetic organization of the gene.

#### Inverse PCR

Inverse PCR was performed as described in my previous study (Watanabe et al., 2018) to obtain the genomic DNA sequence of the coding exons missing in the published genome assembly (Gbim_1.0). Inverse PCRs were repeatedly conducted to obtain the genomic DNA sequence containing the exon(s) encoding the N-terminal region of the *Gryllus* Dsx protein (exon 1 in Figure 1A). Next, the genomic DNA fragment containing the missing coding exon(s) and the upstream sequence of the *Gryllus dsx* gene was amplified.

**Figure 1.**
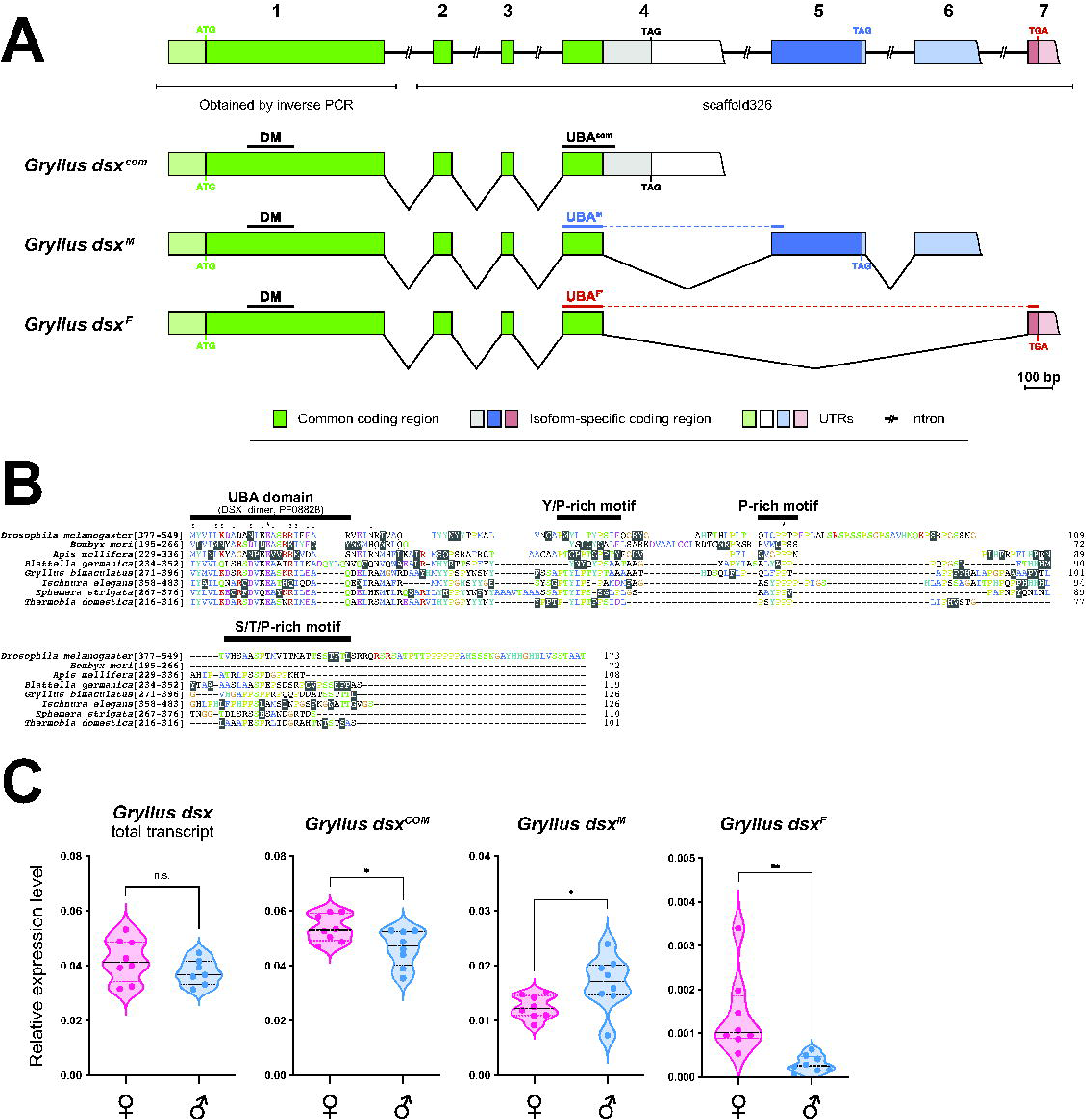
Identification of the *Gryllus dsx* isoforms expressed in the cricket brain. (A) Gene structure of the *Gryllus dsx* gene. The splicing patterns of the three *dsx* isoforms expressed in the brain are illustrated. The *Gryllus dsx* gene consists of seven exons. The genome sequence of exon 1 was obtained in this study, while sequence information for exons 2-7 is available in the genome assembly (scaffold326; GenBank ID: BOPP01000326). The positions of the start and stop codons, as well as the regions encoding the functional domains, are illustrated. (B) Amino acid sequence alignment of the isoform-specific region of the *Gryllus* Dsx^M^ isoform and the corresponding Dsx isoforms from other insects. The position of the C-terminal region of the UBA domain (DSX_dimer, PF08828) and three moderately conserved motifs enriched with Pro, Tyr, Thr, and Ser are indicated. Sequence ranges are indicated with the species name. GenBank IDs: *Drosophila melanogaster*, NP_731197; *Bombyx mori*, NP_001104815; *Apis mellifera*, ABV55178; *Blattella germanica*, QGB21104; *Gryllus bimaculatus*, LC841922; *Ischnura elegans*, XP_046401713; *Ephemera strigata*, LC841917; *Thermobia domestica*, UTQ10360. (C) The expression analysis of the Gryllus dsx total transcripts and isoforms in the brain of adult crickets (n=8, respectively). Dashed lines in the violin plots indicate the 25th to 75th percentile ranges and the central values, respectively. Asterisks indicate statistical significance between the sexes (unpaired t-test; *, p<0.05; **, p<0.01; n.s., p≥0.05).

#### Determination of the transcription start sites (TSSs)

5 ‘RACE was performed to determine the 5 ‘ends of *Gryllus dsx* transcripts. The experimental procedure is described above. The nucleotide sequences of the 5 ‘RACE products were aligned with the genomic DNA sequence obtained previously to map the TSS(s) of the gene.

All PCRs were performed using the Q5 High-Fidelity DNA polymerase. Amplified DNA fragments were cloned into the pGEM-T Easy vector. The nucleotide sequence of the obtained genomic DNA fragment was registered in GenBank (GenBank ID: LC841925). Primers used for genomic DNA amplification are listed in Table 1-1.

### 4. Quantification of the expression levels of *Gryllus doublesex* isoforms in the cricket brain

#### Animals

Male and female adult crickets 7-day after adult emergence were used for gene expression analysis. Crickets were anesthetized on ice, and the brain (the supraesophageal and suboesophageal ganglia without the optic lobes) was dissected in ice-cold phosphate buffered saline (PBS; P-5493, Sigma-Aldrich, Tokyo, Japan), individually homogenized in 100 µl Trizol reagent (Life Technologies, CA, USA), and stored at –70°C until use.

#### RNA extraction and reverse transcription

The reverse transcription for RT-qPCR analysis was conducted following the methods of my previous report (Watanabe et al., 2018).

#### Quantitive PCR (qPCR)

qPCR analysis was performed using the KAPA SYBR FAST qPCR Kit (Kapa Biosystems, MA, USA) and the PCRmax Eco 48 Real-Time qPCR System (PCRmax, Staffordshire, UK), following the methods of my previous report (Watanabe et al., 2018). Gene expression levels were measured by the standard curve method using the EcoStudy Software (ver. 5.0; PCRmax). Ten-fold serial dilutions of standard plasmids containing cDNA fragment(s) of target gene(s) were used to plot standard curves.

Expression levels of target genes/transcripts were normalized with that of the *Gryllus elongation factor 1 alpha* (*ef1*_α_) gene, which showed the highest expression stability in the adult cricket brain (Watanabe et al., 2018). The standard plasmid for *Gryllus ef1*_α_ contained the full-length coding sequence of the gene (GenBank ID: AB583232.1), which was inserted into the pGEM-T Easy vector. The PCR reaction was performed at 50°C for 2 min and 95°C for 5 min, followed by 40 cycles of 95°C for 10 sec and 60°C for 30 sec each. An unpaired t-test was performed to compare the mRNA levels of each *Gryllus dsx* isoform between sexes using the GraphPad Prism for Mac (ver. 10.0; GraphPad Software, MA, USA). The nucleotide sequences of the primers used for RT-qPCR are listed in Table 1-1.

### 5. Sequence comparison and structural analysis

The deduced amino acid sequences of Dsx proteins were aligned using the MAFFT algorithm (Katoh & Standley, 2013) and refined by manual inspection on the Geneious Prime program (ver. 2024.0.4; https://www.geneious.com). The DM domain and the UBA domain of *Gryllus* Dsx protein were annotated using the InterProScan program (Jones et al., 2014; https://www.ebi.ac.uk/interpro/). The alignment was visualized using the ClustalX program (ver. 2.0; Larkin et al., 2007). The helical wheel diagram was drawn using the pepwheel program (https://www.bioinformatics.nl/cgi-bin/emboss/pepwheel). The helical content of the major groove recognition helix of the DM domain at 288K was calculated using the Agadir algorithm (Lacroix et al., 1998; http://agadir.crg.es/about.jsp).

The amino acid sequences of newly identified *dsx* homologs from various insects were included in the sequence comparison. Full-length or partial cDNA sequences of *dsx* homologs were identified in *Gampsocleis buergeri*, *Diestrammena* sp., *Gryllotalpa orientalis*, and *Ephemera strigata* through RT-PCR-based cloning. Illumina short-read RNA-seq was performed in *Ornebius bimaculatus*, *Vescelia pieli ryukyuensis*, *Meloimorpha japonica*, *Oecanthus longicauda*, and *Truljalia hibinonis* to obtain RNA-seq reads encoding *dsx* homologs. Subsequently, RT-PCR-based cloning was conducted to obtain partial cDNA sequences of *dsx* homologs in *Ornebius bimaculatus*, *Meloimorpha japonica*, and *Truljalia hibinonis*. The experimental procedures of RT-PCR-based cloning followed those used for *Gryllus bimaculatus*, while those of the Illumina short-read RNA-seq are described below. RNA extracted from the brain or head of adult males was used for RT-PCR-based cloning and Illumina short-read RNA-seq. The nucleotide sequences of the obtained cDNAs were registered in GenBank (GenBank IDs: *Diestrammena* sp., LC841915; *Gryllotalpa orientalis*, LC841919; *Gampsocleis buergeri*, LC841918; *Ornebius bimaculatus*, LC841924; *Meloimorpha japonica*, LC841923; *Truljalia hibinonis*, LC841926; *Ephemera strigata*, LC841916 and LC841917). Detailed information on the sequences used for the sequence comparison is provided in Table 2.

#### RNA-seq and de novo transcriptome sequence assembly

Total RNA was extracted using the TRIzol reagent (Life Technologies), then purified using the RNeasy Mini Kit (Qiagen, Tokyo, Japan). Genomic DNA contamination was removed by on-column DNase I treatment (RNase-Free DNase Set; Qiagen). RNA samples were submitted to the Rhelixa Next Generation Sequencing service (Tokyo, Japan) for high throughput sequencing. Libraries were generated using the NEBNext Poly(A) mRNA Magnetic Isolation Module and the NEBNext Ultra II Directional RNA Library Prep Kit (New England Biolabs). Sequencing was performed in the NovaSeq 6000 System (Illumina, CA, USA). 123FASTQ (ver. 1.3; Eidi et al., 2024) and SortMeRNA (ver. 4.3.4; Kopylova et al., 2012) were used to remove the Illumina adapter sequences and filter out ribosomal RNA fragments. Then, *de novo* sequence assembly was conducted using the Trinity assembler (v2.14.0; Grabherr et al., 2011). The RNA-seq raw reads were registered in BioProject (BioProject ID: PRJDB18769).

### 7. Systematic Evolution of Ligands by Exponential enrichment (SELEX)

#### Bait protein preparation

The DM domain of the Dsx proteins from various insect and crustacean species (*Drosophila melanogaster*, *Apis mellifera*, *Halyomorpha halys*, *Gryllus bimaculatus*, *Ephemera strigata*, *Thermobia domestica*, and *Daphnia pulex*) was subjected to SELEX. Since *Daphnia pulex* has two *dsx* homologs, *dsx1*, which plays a role in sex determination, was included in this analysis. Considering that the *Gryllus bimaculatus* Dsx protein contains a highly substituted recognition helix, I constructed a “back-mutated” version of the protein, incorporating the recognition helix from *Gampsocleis buergeri*, to use as a bait protein (*Gryllus bimaculatus* DsxDM^BM^). The N-terminal region of the Dsx proteins, N-terminally fused to a Twin-Strep tag (WSHPQFEK-GGGSGGGSGG-SAWSHPQFEK) and a 2xGS linker (GGGGS-GGGGS), was used as the bait protein (Figure 4A).

**Figure 4.**
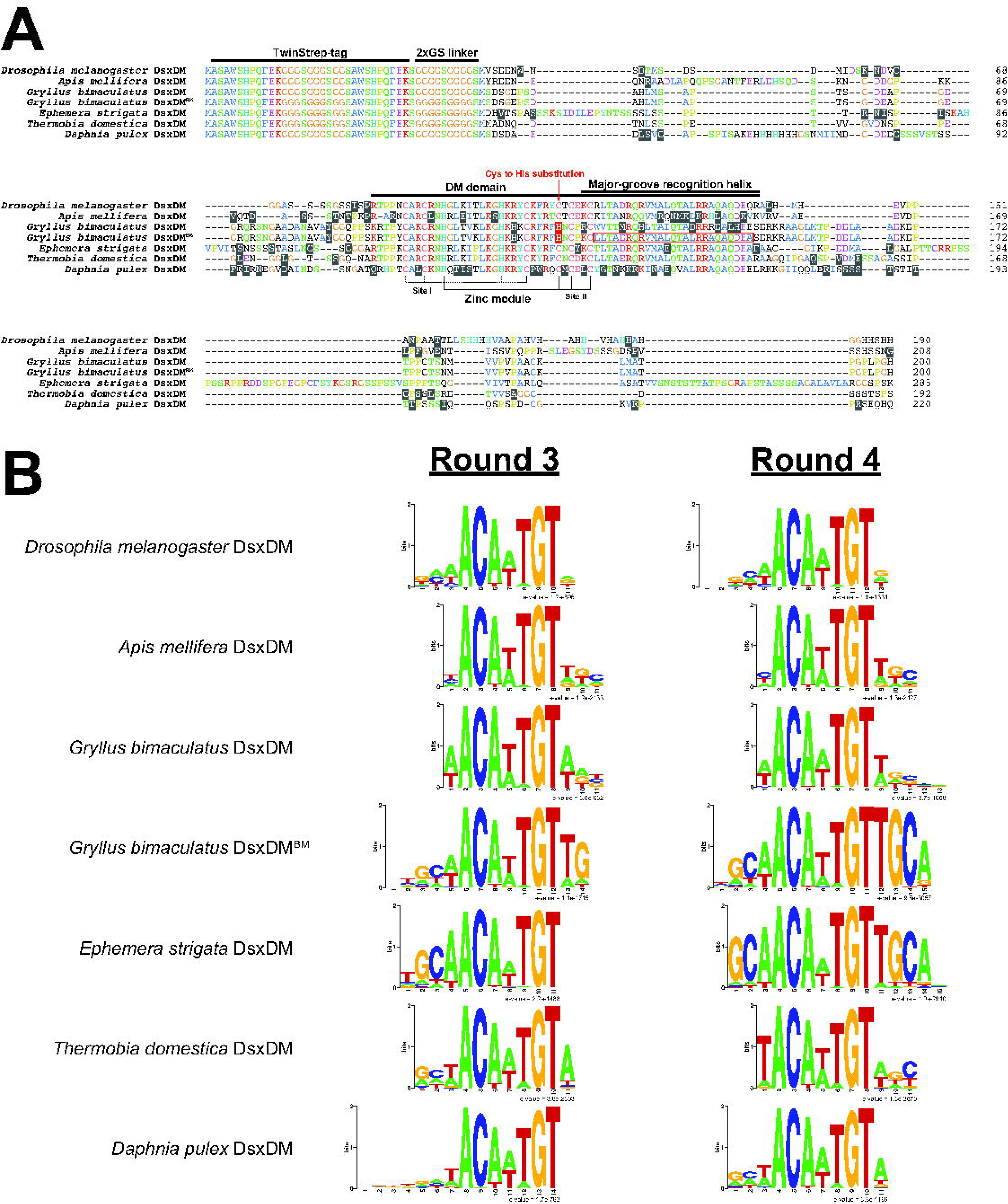
The target DNA sequences of the DM domain of Dsx proteins of various insect and crustacean species revealed by SELEX. (A) Amino acid sequence alignment of the seven bait proteins used in the SELEX experiment. All proteins contain a TwinStrep-tag and a 2xGS linker at their N-termini. The position of the DM domain and residues critical for DM domain conformation and function are highlighted. The ‘back-mutated ‘recognition helix in the *Gryllus bimaculatus* DsxDM^BM^ is outlined in red. (B) Sequence logos represent the motifs identified in the SELEX libraries from rounds 3 and 4. A random DNA library containing a 26-bp random sequence was selected using seven DM domains representing insect phylogeny and sequence variation among Dsx homologs. A 7-nt palindromic motif, ACAyTGT, was identified with all seven DM domains.

The coding sequences of the DM domain of insect Dsx proteins were amplified by PCR from the cDNA libraries prepared in this study or those from my previous work (Watanabe et al., 2010). Forward and reverse primers with BamHI and EcoRI sites attached to their 5 ‘ends, respectively, were used for RT-PCR. The coding sequence of the N-terminal region of *Daphnia pulex* Dsx1 protein (a.a. 1-177 of BAM33607; *Daphnia pulex* DsxDM) was chemically synthesized with the coding sequence of the Twin-Strep tag and 2xGS linker (gBlock; Integrated DNA Technologies, IA, USA). A BamHI site was placed between the 2xGS linker coding sequence and the DM domain, while an EcoRI site was positioned 3 ‘adjacent to the stop codon. The NEBuilder HiFi DNA Assembly Master Mix (New England Biolabs) was used to integrate the coding sequence of the Twin-Strep-tagged *Daphnia pulex* DsxDM into EcoRV/EcoRI-digested pTD1 plasmid (Shimadzu, Shiga, Japan). Subsequently, the plasmid was digested with BamHI and EcoRI, and the *Daphnia pulex* DsxDM coding sequence was replaced with the coding sequence of the DM domain from other insect species.

The mRNAs for cell-free protein expression were transcribed using the HiScribe T7 Quick High Yield RNA Synthesis Kit (New England Biolabs) according to the manufacturer’s instructions. A 1 µg template DNA was transcribed in a 20 µl reaction. The cell-free protein expression was performed using the Transdirect Insect Cell system (Shimadzu) according to the manufacturer’s instructions. A total of 16 µg of RNA was added to the 50 µl reaction. After a 5-hour incubation at 25°C, the reaction was aliquoted, flash-frozen in liquid nitrogen, and stored at – 70°C.

#### Random library preparation

Library DNA containing a 26-bp random sequence was prepared according to Huang et al. (1993) with modifications. A pool of single-stranded random oligonucleotides (oMC070m; >Table 1-2) with a 26-bp variable sequence was chemically synthesized (Integrated DNA Technologies). oMC070m and oMC069 (Table 1-2) were used to convert the single-stranded random library into a double-stranded DNA library using the Klenow fragment (New England Biolabs). After treatment with Exonuclease I to remove ssDNA, the dsDNA library was purified by phenol/chloroform extraction followed by isopropanol precipitation. Then, the dsDNA library was amplified by 15-cycle PCR using the Q5 High-Fidelity DNA Polymerase.

**Table 1-2.**
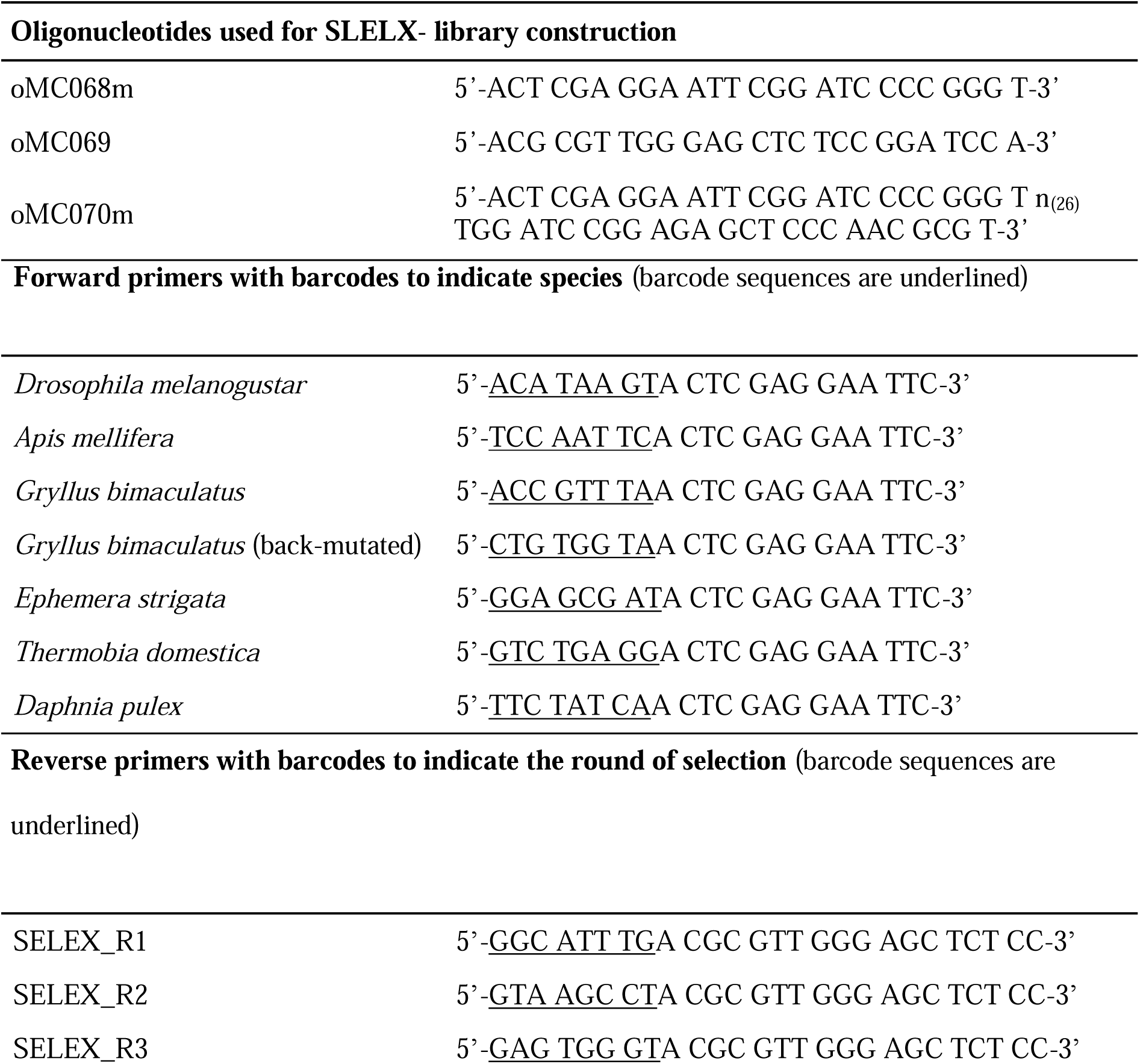
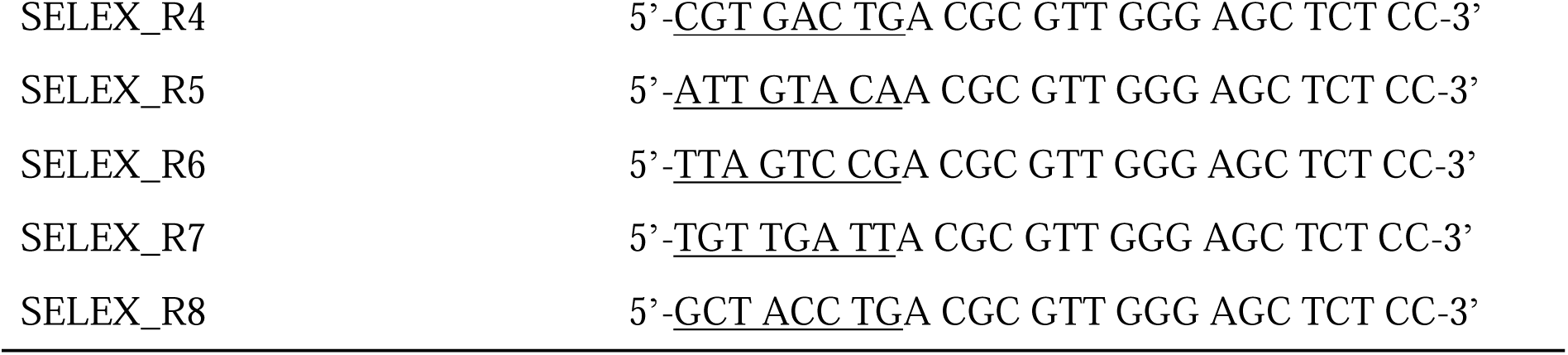
List of primers used for SLELX-library construction and barcoding.

#### SELEX

25 µl suspension of MagStrep “type3” XT beads (IBA Lifesciences, Göttingen, Germany) were equilibrated twice with 250 µl of EMSA buffer (20 mM HEPES, pH 8.0; 100 mM KCl; 20% glycerol; 5 mM MgClL). Then, 4 µl of the bait protein solution, the equilibrated beads, and the SELEX library were diluted in 100 µl of EMSA buffer containing 1 mM dithiothreitol, 100 mM zinc acetate, 25 ng/µl Herring sperm DNA, 50 ng/µl poly(dI-dC)•poly(dI-dC), 1 µg/µl BSA, and the protease inhibitor cocktail (S8830-2TAB; Sigma-Aldrich). In the first and second rounds of SELEX, 200 ng of library DNA was added; from round 3 onwards, 1 ng of library DNA was added to the reaction. The mixture was incubated for 20 minutes at 25°C with gentle shaking. The beads were then collected using a magnetic rack, washed five times with 100 µl of EMSA buffer, and dissolved in 10 µl of TE (pH 8.0).

The selected libraries were amplified by PCR using the Q5 High-Fidelity DNA Polymerase (New England Biolabs), with oligonucleotides oMC068m and oMC069 as primers (Table 1-2). A 2.5 µl suspension of the beads was added to the 50 µl PCR reaction. The selected libraries from the first round of SELEX were amplified through a 20-cycle PCR; from the second round onwards, the libraries were amplified through a 15-cycle PCR. Amplified libraries were concentrated by isopropanol precipitation, dissolved in TE (pH 8.0), and stored at –20°C.

#### Barcoding and Sequencing of the SELEX library

Barcoded SELEX libraries were produced by PCR amplification with primers containing barcode sequences. The forward primers contain barcodes to discriminate between species, while the reverse primers contain barcodes to indicate the round of selection (Table 1-2). After purification by phenol/chloroform extraction followed by isopropanol precipitation, equal amounts of the barcoded SELEX libraries were mixed and submitted to the Hokkaido System Science (Sapporo, Japan) for high throughput sequencing.

Libraries were generated using the NEBNext Ultra II DNA Library Prep Kit (PCR Free Library) (New England Biolabs). Sequencing was performed in the NovaSeq 6000 System (Illumina).

#### Data analysis

After dividing the data by species and rounds based on the barcode sequences on the Geneious Prime program, representative sequence motifs were identified using the MEME software (ver. 5.4.1; Bailey & Elkan, 1994).

### 8. Quantification of gene copy numbers of *dsx* homologs and other genes in orthopteran insects

To determine whether the genes of interest (GOIs) are located on an autosome or the X chromosome, their copy numbers were measured by qPCR and compared between the sexes. Genomic DNA was extracted from *Acheta domestica*, *Gampsocleis buergeri*, *Locusta migratoria*, and *Teleogryllus commodus* using the Wizard Genomic DNA Purification Kit (Promega).

#### qPCR

The copy numbers of these GOIs were quantified by qPCR. qPCR analysis was performed as described above. Gene expression levels were measured by the ΔΔCt method. The *glucose-6-phosphate dehydrogenase* (*G6PD*) gene, which was reported as an X-linked gene in mole cricket (Rao & Padmaja, 1992), was used as a positive control for the X-linked gene. *Odorant receptor co-receptor* (*ORco)* and *ef1*α genes were used as internal reference genes. *ORco* gene is reported to be autosomal in *Locusta migratoria* (Li et al., 2016). For *Gryllus bimaculatus*, a genomic region known to be autosomal, where the transgene was integrated in my previous study (line#19; Watanabe et al., 2018), was used as an additional internal standard. The copy number of each GOI was measured in males and females (n=4, respectively), and normalized to the geometric mean of the internal reference genes. An unpaired t-test was performed to compare the copy number of each GOI between sexes using the GraphPad Prism version 10.0 for Mac (GraphPad Software).

#### Selection of GOIs

To obtain genomic DNA or cDNA fragments encoding the *ORco*, *ef1*α, and *G6PD* homologs in five orthopteran insect species, tBLASTn searches were performed on the genomic scaffolds of *Locusta migratoria*, *Gryllus bimaculatus*, and *Acheta domesticus* (BioProject IDs: PRJNA185471, PRJDB10609, PRJNA612585, and PRJNA706033, respectively), as well as on the TSA of *Gampsocleis buergeri* and *Teleogryllus commodus* (BioProject IDs: PRJNA352290 and PRJNA252786, respectively), with the deduced amino acid sequences of these selected genes, GOIs. In addition to the genes described above, a representative set of genes on the X chromosome and chromosome 1 of the *Schistocerca americana* genome (iqSchAmer2.1; BioProject ID: PRJNA772266) were selected as GOIs. With the deduced amino acid sequences of these selected genes, tBLASTn searches were performed on the genomic scaffolds of *Gryllus bimaculatus* and *Locusta migratoria*. The nucleotide sequences of the primers used for qPCR are listed in Table 1-3.

**Table 1-3.**
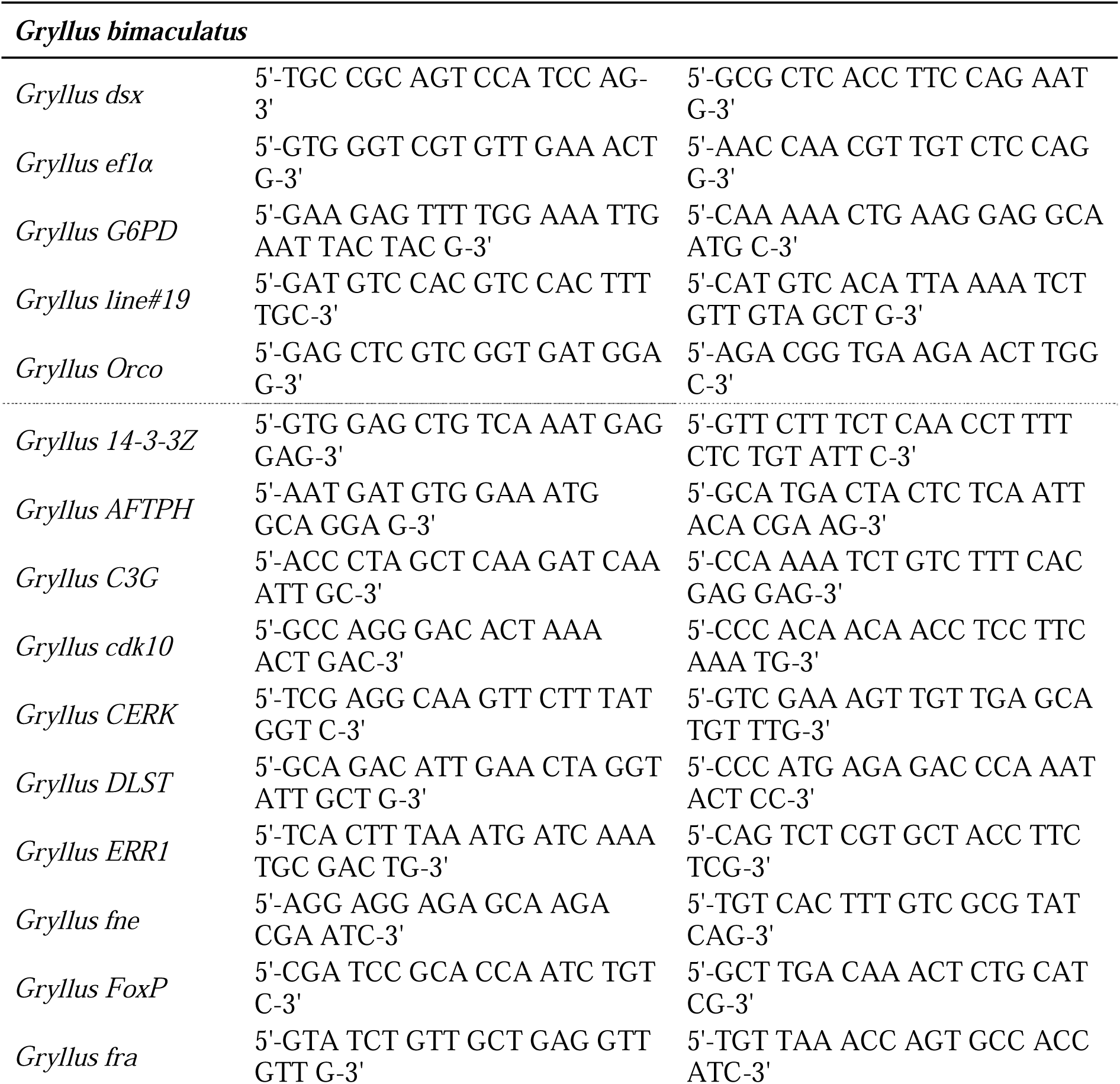
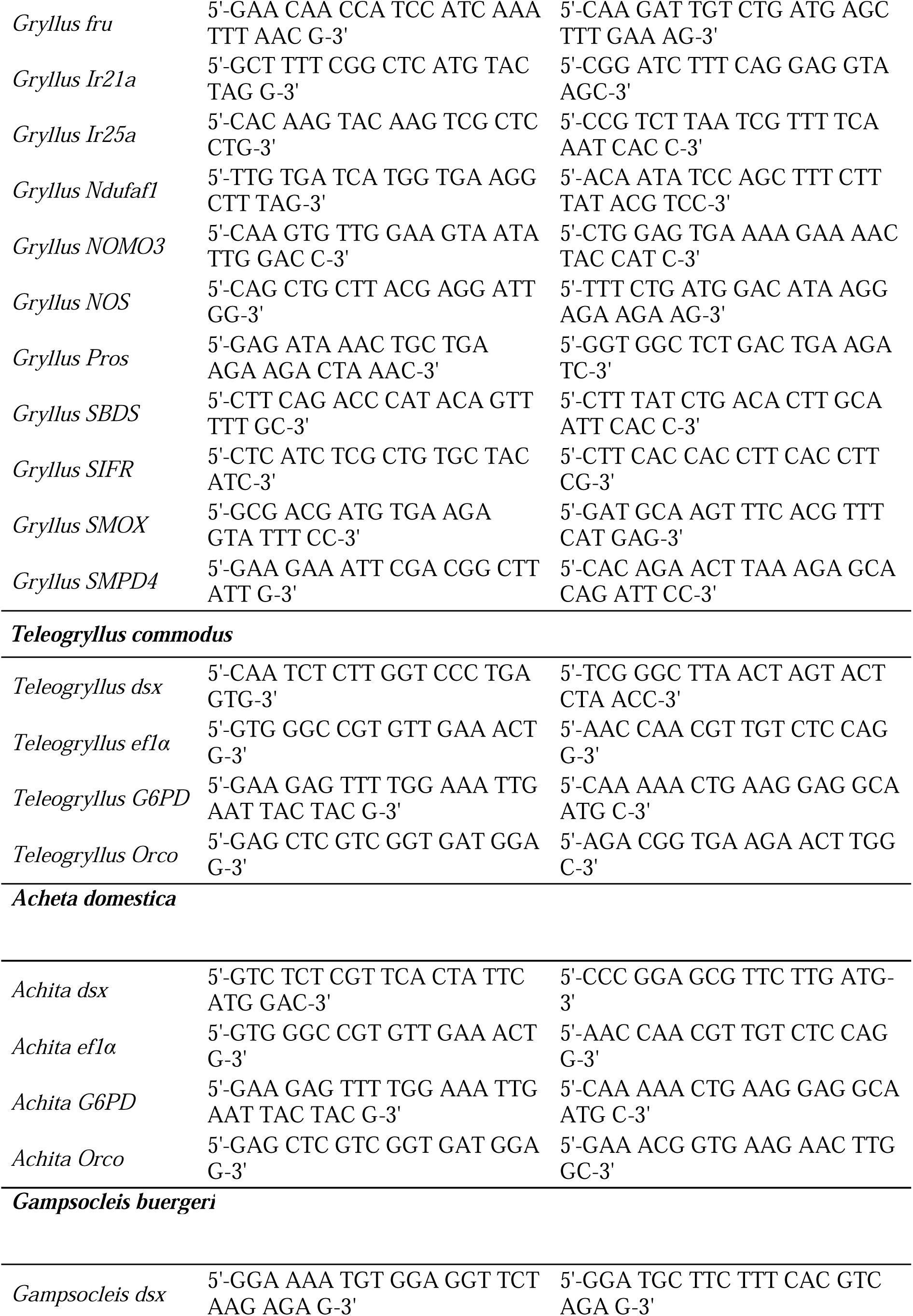
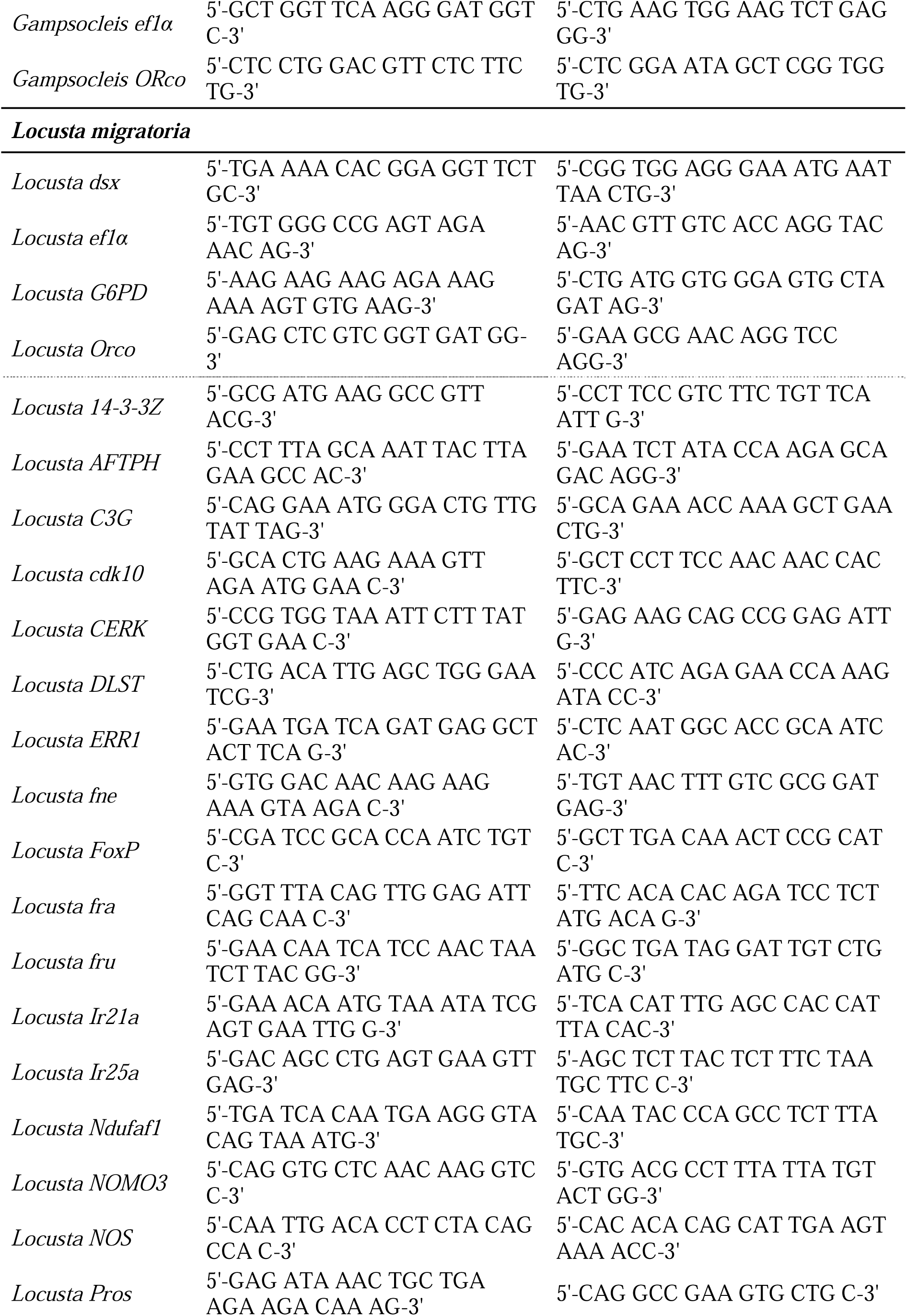
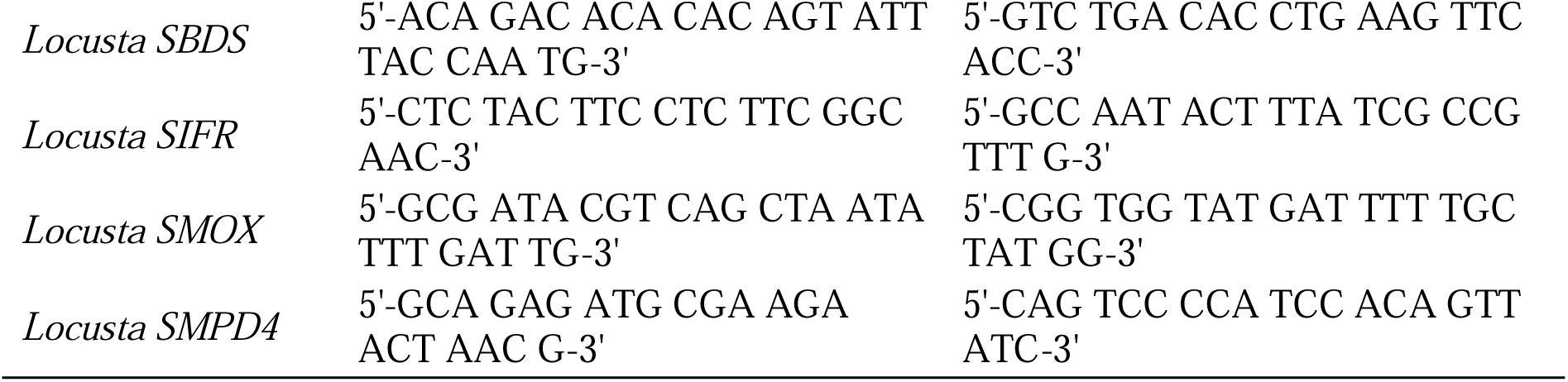
List of primers used for the gene copy number analysis.

### 9. Generation of *Gryllus dsx* deletion mutant

#### Design of CRISPR RNA (crRNA)

Guide RNAs (gRNAs) were designed to introduce indels in the DM domain of the *Gryllus* Dsx protein. On-target cleavage efficiency was predicted using the DeepSpCas9 web tool (Kim et al., 2019; https://deepcrispr.info/DeepSpCas9/). Off-target scores against the *Gryllus bimaculatus* genome assembly (Gbim_1.0) were calculated using the Geneious Prime program. The crRNA was chemically synthesized (CRISPR-Cas9 crRNA; Integrated DNA Technologies) and annealed with the CRISPR-Cas9 tracrRNA (Integrated DNA Technologies) to form the gRNA heteroduplex, following the manufacturer’s instructions.

#### Microinjection

The Alt-R S.p. Cas9 Nuclease V3 (Integrated DNA Technologies) and gRNA heteroduplex were dissolved in the Nuclease-Free Duplex Buffer (Integrated DNA Technologies) at final concentrations of 1.2 µM and 1.35 µM, respectively, and incubated at room temperature for 30 minutes to form Cas9 ribonucleoprotein complex (RNP). The Cas9 RNP solution was injected into fertilized eggs of the Hokudai g*white* strain. Eggs were laid in wet sand and collected 30-60 min within egg laying. They were rinsed with 70% ethanol, aligned on the hand-made injection chamber, and kept on ice. Micropipettes were pulled on a micropipette puller (P-2000 Laser-Based Micropipette Puller; Sutter Instrument Co., Novato, CA, USA) using thin-walled filament glass capillary (GDC-1; Narishige). The capillaries were backfilled with the injection solution using the GELoader tip (Eppendorf, Tokyo, Japan) and connected to the IM-400 microinjector (Narishige, Tokyo, Japan). Cas9 RNP solution was injected into the dorsal posterior part of the eggs. Eggs were incubated at 28°C until hatching.

#### Genetics

Cas9 RNP-injected embryos were incubated at 28°C until hatching. The crickets were raised to adulthood and inbred. Female G1 crickets were raised to adulthood and backcrossed to the Hokudai *gwhite* strain. After collecting fertilized eggs, the G1 crickets were sacrificed, and genomic DNA was extracted from their hind legs using the Wizard Genomic DNA Purification Kit (Promega).

#### Selection

G1 crickets harboring indels were selected using the T7E1 assay. For the T7E1 assay, a ∼700-bp genomic DNA fragment was amplified using the Q5 High-Fidelity DNA Polymerase (New England Biolabs) and subjected to the T7E1 assay with the Alt-R Genome Editing Detection Kit (Integrated DNA Technologies) according to the manufacturer’s instructions. Subsequently, the genomic region encoding the DM domain was PCR amplified again in individuals confirmed not to carry the wild-type allele, and the sequence was determined to verify the presence of the InDel.

Females with InDel were crossed with the Hokudai *gwhite* males. Primers used for the T7E1 assay are listed in Table 1-4.

**Table 1-4.**
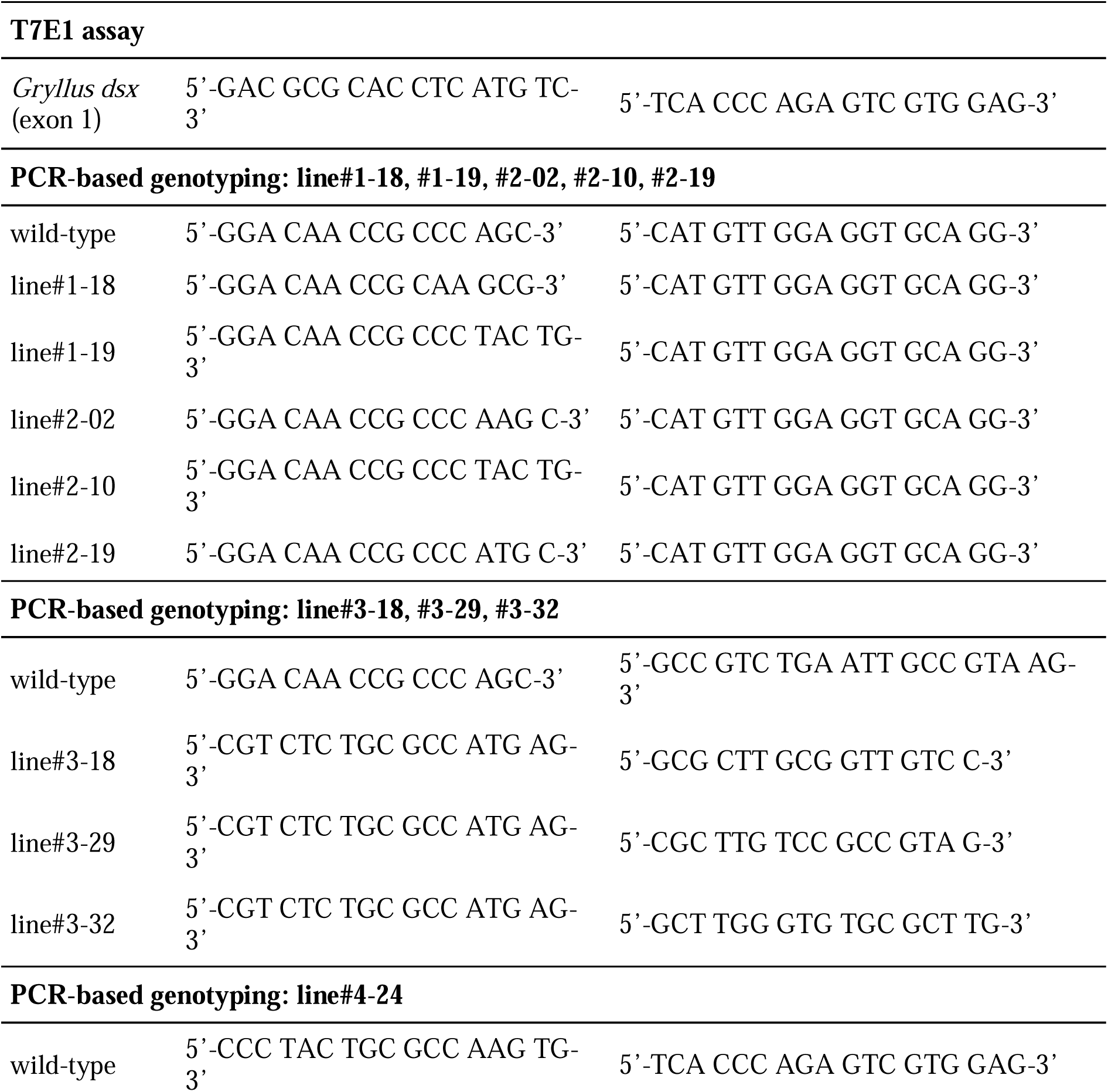
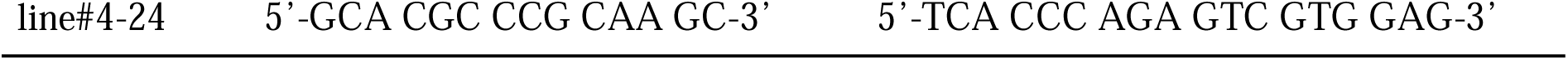
List of primers used for generation of knockout lines.

#### PCR-based genotyping

Genomic DNA was extracted from the tip of a hindwing, or hind legs using the Wizard Genomic DNA Purification Kit and dissolved in 25 µl of TE buffer (pH 8.0). PCR was performed using the OneTaq DNA Polymerase (New England Biolabs) in a 5 µl reaction, with 0.3 µl of genomic DNA solution added. For genotyping, forward and reverse primers were designed to distinguish between the wild-type allele and the deletion allele (Table 1-4). PCR products were resolved on a 1.5% agarose gel in TBE buffer and visualized using ethidium bromide.

### 10. Behavioral experiment

Adult crickets, one week after the imaginal molt, were used for behavioral experiments. Crickets exhibiting female morphology were collected from a colony of *Gryllus dsx*^del^ mutants. A wild-type cricket (Hokudai WT strain) was used as an interaction partner. On the day of the imaginal molt, crickets were housed in separate cases according to sex to prevent mating. Prior to the behavioral experiment, each cricket was individually isolated in a 100 ml beaker (ø4.5 cm) for three days. A piece of mouse food was placed in the beaker, and water was supplied in a 2 ml tube with a cotton plug. A pair of crickets was introduced into an acrylic case (15 cm x 15 cm x 15 cm) with paper towels seated on the bottom. We observed their behavior for over 5 minutes, and their interactions were recorded using an iPhone 11 (Apple, CA, USA). The behavior of the crickets collected from a colony of *Gryllus dsx*^del^ mutants was classified according to the criteria in Table 3, based on Adamo & Hoy (1994). After the behavioral experiment, the hind legs were dissected and stored in 70% ethanol for PCR-based genotyping. The experimental procedure for PCR-based genotyping is described above.

**Table 2.** List of Dsx homologs used for the comparison of the amino acid sequence in the DM domain.

| Species | Protein |  | GenBank ID | Sequence range |
| --- | --- | --- | --- | --- |
| *Drosophila melanogaster* | Dsx | Protein | NP_731197 | a.a. 64-98 |
| *Bactrocera dorsalis* | Dsx | Protein | XP_011203913 | a.a. 52-86 |
| *Ceratitis capitata* | Dsx | Protein | AAN63597 | a.a. 64-98 |
| *Anastrepha amita* | Dsx | Protein | ABF50951 | a.a. 64-98 |
| *Aedes aegypti* | Dsx | Protein | ABD96573 | a.a. 58-92 |
| *Bombyx mori* | Dsx | Protein | NP_001104815 | a.a. 55-89 |
| *Danaus plexippus* | Dsx | Protein | EHJ78146 | a.a. 55-89 |
| *Antheraea mylitta* | Dsx | Protein | ADL40853 | a.a. 55-89 |
| *Helicoverpa armigera* | Dsx | Protein | AHF81649 | a.a. 55-89 |
| *Ostrinia scapulalis* | Dsx | Protein | BAJ25851 | a.a. 55-89 |
| *Annulipalpia sp.* | Dsx | TSA | GATX01084595 | bases 890-786 |
| *Boreus hyemalis* | Dsx | TSA | GAYK01000638 | bases 363-467 |
| *Oropsylla silantiewi* | Dsx | TSA | GAWY01012876 | bases 700-804 |
| *Apis mellifera* | Dsx | Protein | NP_001104725 | a.a. 82-116 |
| *Harpegnathos saltator* | Dsx | Protein | EFN87725 | a.a. 50-84 |
| *Cerapachys biroi* | Dsx | Protein | EZA59919 | a.a. 50-84 |
| *Fopius arisanus* | Dsx | Protein | JAG85033 | a.a. 52-86 |
| *Microplitis demolitor* | Dsx | Protein | XP_008554550 | a.a. 52-86 |
| *Trichomalopsis sarcophagae* | Dsx | Protein | ACJ65503 | a.a. 53-87 |
| *Nasonia vitripennis* | Dsx | Protein | NP_001155990 | a.a. 53-87 |
| *Athalia rosae* | Dsx | Protein | XP_012262273 | a.a. 51-85 |
| *Tribolium castaneum* | Dsx | Protein | EFA07556 | a.a. 49-83 |
| *Trypoxylus dichotomus* | Dsx | Protein | BAM 93346 | a.a. 48-82 |
| *Onthophagus sagittarius* | Dsx | Protein | AEX92944 | a.a. 48-82 |
| *Osmylus fulvicephalus* | Dsx | TSA | GAYC01086997 | bases 1,073-969 |
| *Conwentzia psociformis* | Dsx | TSA | GAYH01007785 | bases 480-584 |
| *Corydalus cornutus* | Dsx | TSA | GATG01001294 | bases 960-856 |
| *Inocellia crassicornis* | Dsx | TSA | GAZH01013937 | bases 972-868 |
| *Halyomorpha halys* | Dsx | Protein | XP_014275563 | a.a. 57-91 |
| *Cimex lectularius* | Dsx | Protein | XP_014246101 | a.a. 57-91 |
| *Acyrtosiphon pisum* | Dsx | Protein | XP_001949339 | a.a. 62-96 |
| *Diaphorina citri* | Dsx | Protein | XP_017303801 | a.a. 34-68 |
| *Bemisia tabaci* | Dsx | Protein | XP_022184822 | a.a. 57-91 |
| *Nilaparvata lugens* | Dsx | Protein | XP_018913623 | a.a. 60-94 |
| *Okanagana villosa* | Dsx | Protein | XP_002427670 | a.a. 65-99 |
| *Pediculus humanus corporis* | Dsx | TSA | GAWR01005396 | bases 1,429-1,325 |
| *Menopon gallinae* | Dsx | TSA | GAYV01012131 | bases 468-364 |
| *Liposcelis bostrychophila* | Dsx | TSA | GAWQ02048299 | bases 469-573 |
| *Mantis religiosa* | Dsx | TSA | GASW01220805 | bases 400-504 |
| *Metallyticus splendidus* | Dsx | TSA | GATB01322676 | bases 185-289 |
| *Blattella germanica* | Dsx | Protein | QGB21104 | a.a. 57-91 |
| *Areataon asperrimus* | Dsx | TSA | GAWC01128745 | bases 197-301 |
| *Locusta migratoria* | Dsx | TSA | AVCP011201136 | bases 115-219 |
| *Tetrix subulata* | Dsx | TSA | GASQ01124537 | bases 106-210 |
| *Gampsocleis buergeri* | Dsx | Protein | LC841918 | a.a. 25-59 |
| *Diestrammena sp.* | Dsx | Protein | LC841915 | a.a. 25-59 |
| *Gryllotalpa orientalis* | Dsx | Protein | LC841919 | a.a. 52-86 |
| *Ornebius bimaculatus* | Dsx | Protein | LC841924 | a.a. 61-95 |
| *Vescelia pili ryukyuensis* | Dsx | RNA-seq raw read | A00609:679:HY2VLDSX 5:2:2345:22010:4413 | bases 35-139 |
| *Meloimorpha japonica* | Dsx | Protein | LC841923 | a.a. 36-71 |
| *Laupala kohalensis* | Dsx | Genome | NNCF01126652 | base 651,787-651,894 |
| *Apteronomobius asahinai* | Dsx | TSA | IACC01080379 | bases 812-708 |
| *Oecanthus longicauda* | Dsx | RNA-seq raw read | A00609:936:H73LLDSX C:2:1663:4056:5822 | bases 32-139 |
| *Truljalia hibinonis* | Dsx | Protein | LC841926 | a.a. 84-118 |
| *Teleogryllus commodus* | Dsx | TSA | GBHB01043004 | bases 2,425-2,321 |
| *Gryllus bimaculatus* | Dsx | Protein | LC841921 | a.a. 74-108 |
| *Forficula auricularia* | Dsx | TSA | GAYQ01175103 | bases 137-241 |
| *Epiophlebia superstes* | Dsx | TSA | GAVW02000373 | bases 733-837 |
| *Ichnura elegans* | Dsx | TSA | GFWX01082137 | bases 3,058-2,954 |
| *Ephemera strigata* | Dsx | Protein | LC841916 | a.a. 92-126 |
| *Ecdyonurus insignis* | Dsx | TSA | GCCL01025403 | bases 823-179 |
| *Thermobia domestica* | Dsx | Protein | UTQ10360 | a.a. 66-100 |
| *Holacanthella duospinosa* | Dsx | TSA | GFPE01037288 | bases 1,474-1,370 |
| *Acerentomon sp.* | Dsx | TSA | GAXE01026579 | bases 1,146-1,042 |
| *Daphnia pulex* | Dsx1 | Protein | BAM 33607 | a.a. 94-128 |
| *Ceriodaphnia dubia* | Dsx1 | Protein | BAM 33611 | a.a. 89-123 |
| *Daphnia pulex* | Dsx2 | Protein | BAM 33608 | a.a. 67-101 |
| *Ceriodaphnia dubia* | Dsx2 | Protein | BAM 33612 | a.a. 76-110 |
| *Penaeus monodon* | Dsx | Protein | ACC94178 | a.a. 40-74 |
| *Homo sapiens* | DMRT1 | Protein | NP_068770 | a.a. 96-130 |

**Table 3.**
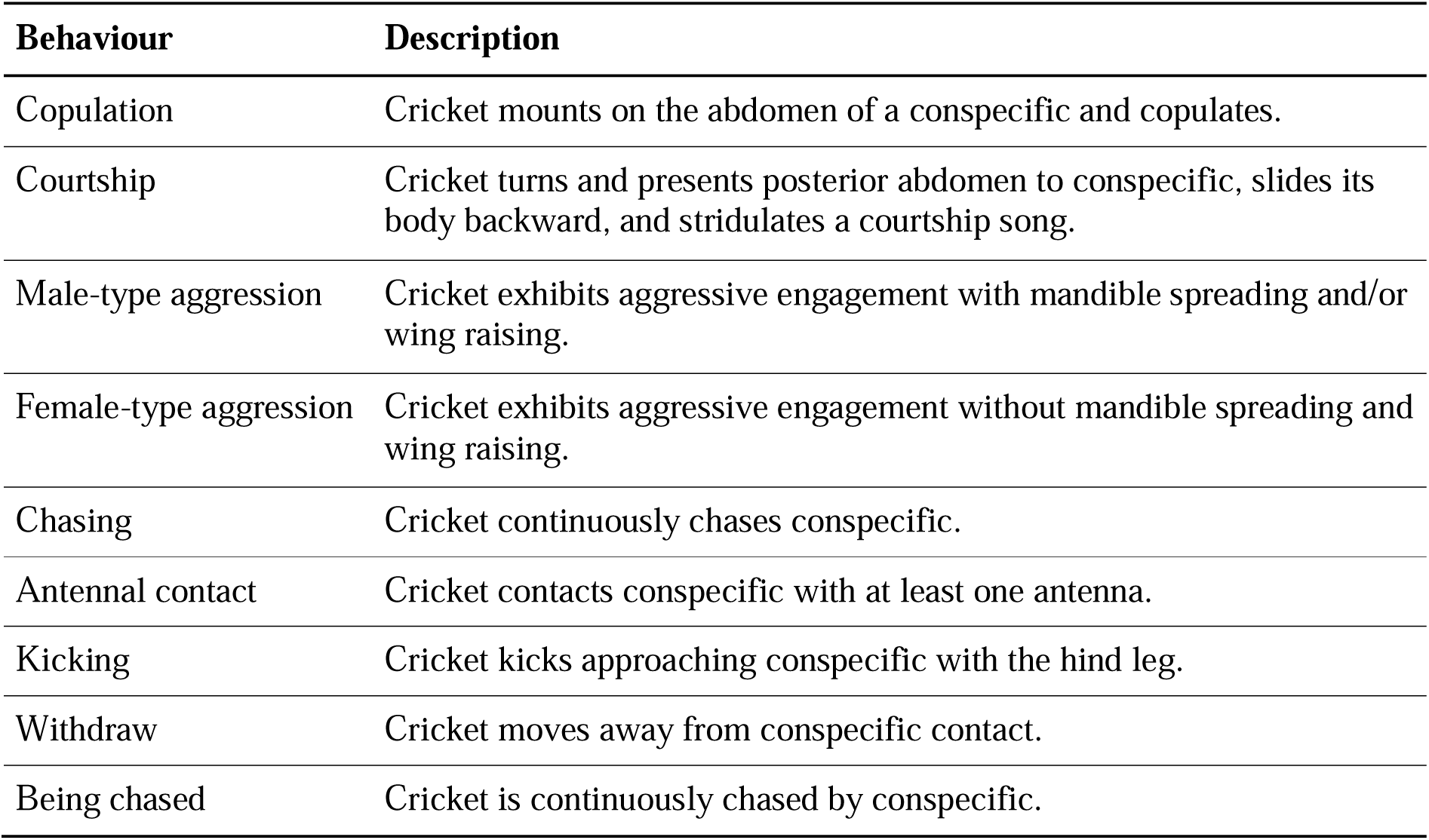
Behavioral categories of social interactions observed between a pair of adult crickets.

## Results

### 1. *Gryllus dsx* isoforms exhibited sex-biased expression in the brain

The brain of the two-spotted cricket, *Gryllus bimaculatus*, expresses three *dsx* isoforms, each encoding Dsx proteins with a set of conserved functional domains (the DM domain (DM DNA-binding domain: IPR001275) and the UBA domain (Doublesex dimerization domain: PF08828); Figure 1A). These three isoforms differ in their C-terminal tails: the *Gryllus dsx^M^* isoform encodes a Dsx protein with the longest isoform-specific C-terminal tail (103 amino acid residues), which resembles the isoform-specific region of male-type Dsx isoforms found in other insect species (Figure 1B). The *Gryllus dsx^F^* and *dsx^COM^* isoforms encode Dsx proteins with shorter isoform-specific C-terminal tails (10 and 54 amino acid residues, respectively), which show no sequence homology to known Dsx isoforms in other insects.

The *Gryllus dsx* gene is composed of seven exons (Figure 1A). The first four exons are shared among the three *Gryllus dsx* isoforms, while exons 5 and 6, which encode the male-specific C-terminal tail, are included only in the *Gryllus dsx^M^* isoform. In the *Gryllus dsx^F^* isoform, exons 5 and 6 are excluded, and exon 4 is spliced directly to exon 7. The *Gryllus dsx^COM^* isoform is generated by intron retention within exon 4, which encodes the common UBA domain. 5 ‘RACE-based TSS analysis revealed that the *Gryllus dsx* gene in the brain is transcribed from a single TSS located 242 bp upstream of the first ATG of the gene.

Since *dsx* homologs show sex-specific expression in other insect species, the expression levels of *Gryllus dsx* isoforms in the brain were compared between adult males and females (Figure 1C). The mRNA expression levels of the total *Gryllus dsx* transcript did not significantly differ between the brains of male and female adults. The *Gryllus dsx^M^* isoform was detected in the brains of both sexes, but its mRNA expression level is significantly higher in male adults compared to female adults. The *Gryllus dsx^F^* isoform was detected in female brains but rarely in male brains, with its mRNA expression level significantly higher in female adults than in male adults. The *Gryllus dsx^COM^* isoform was detected in both female and male brains, with its mRNA expression level significantly higher in female adults than in male adults. These data indicate that the *Gryllus dsx* isoforms are expressed in the brain in a sex-biased manner, rather than the sex-restricted expression observed in *Drosophila* and other insect species.

### 2. A high rate of substitutions was detected in the DM domain of cricket Dsx homologs

A significant number of amino acid substitutions were detected in the *Gryllus* Dsx protein when compared to the *Drosophila* Dsx protein. Notably, the DM domain, which is crucial for DNA-binding function, contains many of these substitutions (Figure 2A). The DM domain includes intertwined CCHC-type and HCCC-type zinc binding sites (sites I and II) necessary for proper conformation, as well as the major-groove recognition helix for base-specific contact (Figure 2A; Zhu et al., 2000). In the *Gryllus* Dsx protein, the second amino acid residue responsible for zinc coordination in the second zinc-binding site (site II) was replaced with His, altering the canonical HCCC-type zinc-binding site to an HHCC-type zinc finger. Additionally, >50% of the sites in the major-groove recognition helix were substituted compared to those in *Drosophila*. The finding that the DM domain of the *Gryllus* Dsx protein exhibits a high degree of substitution was unexpected, given that amino acid sequences in the DNA-binding domains of transcription factors are typically conserved across species (e.g., see Figure 3-1 in Watanabe et al., 2018). However, this finding is based on a pairwise comparison between the *Drosophila* and *Gryllus* Dsx proteins, leaving open the possibility that the DM domain of insect Dsx proteins might naturally be more prone to amino acid substitutions.

**Figure 2.**
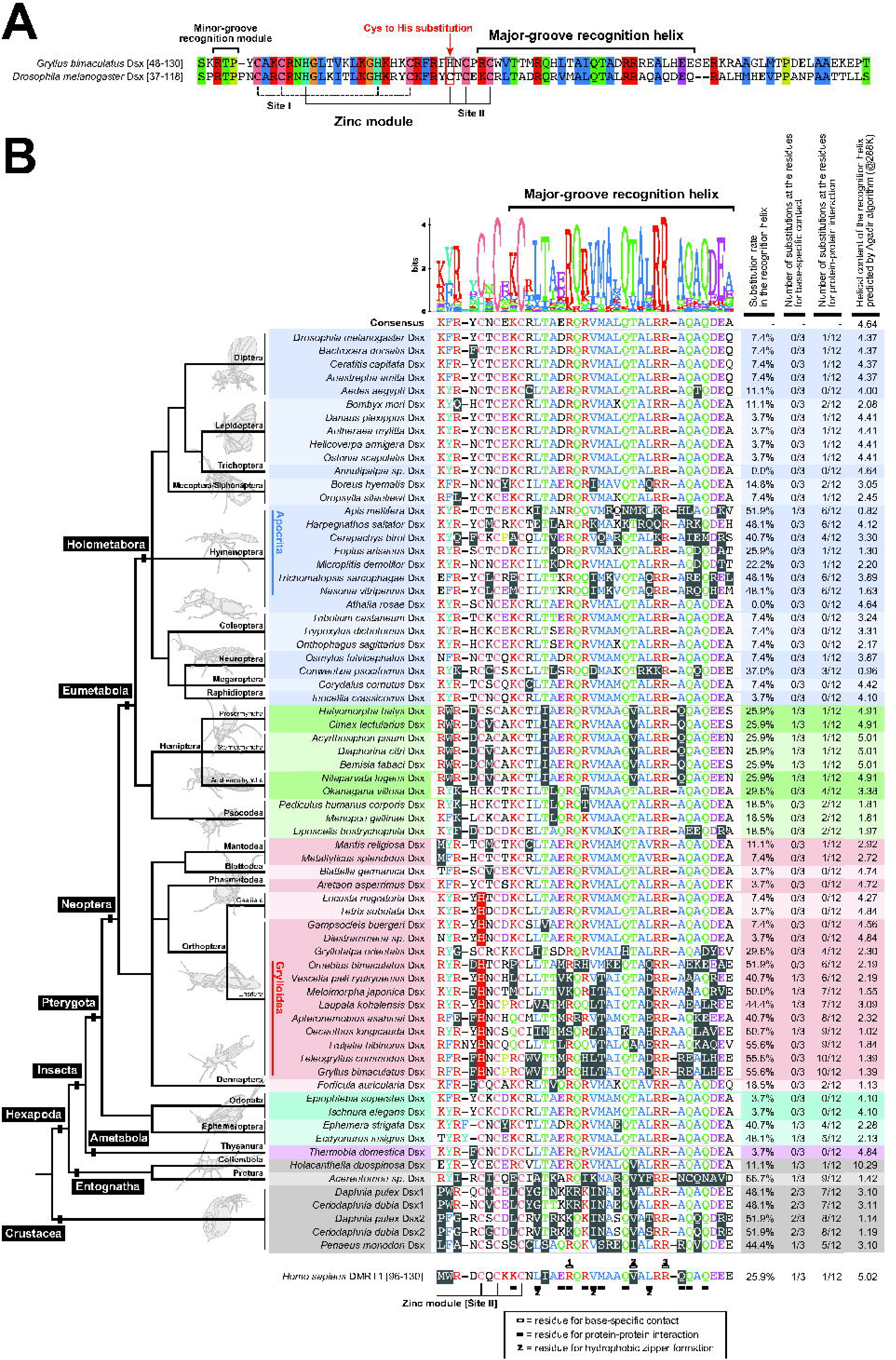
Comprehensive comparison of the amino acid sequence of the DM domain among insect and crustacean Dsx homologs. (A) Pairwise sequence alignment of the DM domain of *Gryllus* Dsx and *Drosophila* Dsx. Positions of the major-groove recognition helix, the minor-groove recognition helix, cysteine and histidine residues coordinating zinc ions are illustrated. Conserved residues are colored according to amino acid type, following the Clustal X color scheme. The position of the Cys to His substitution in the zinc module is illustrated. (B) Multiple sequence alignment of the major-groove recognition helix of various insect and crustacean Dsx proteins. Residues are colored according to amino acid type, following the Clustal X color scheme. Exceptional residues are highlighted as white letters on a gray background. The Cys to His substitutions in the zinc module are highlighted as white letters on a red background. The amino acid sequence of the major-groove recognition helix of *Homo sapiens* DMRT1 protein, along with the positions of the functional residues identified by X-ray crystallography (Murphy, et al., 2015), is shown below the alignment. The phylogenetic tree is positioned to the left of the alignment. The positions of two lineages—the suborder Apocrita (Hymenoptera) and the superfamily Grylloidea (Orthoptera: Ensifera)—whose Dsx homologs possess highly substituted major-groove recognition helices are indicated. The substitution rate, the number of substitutions at residues involved in base-specific contacts, the number of substitutions at residues involved in protein-protein interactions, and the helical content of the recognition helix predicted by the Agadir algorithm are summarized in the table positioned to the right of the alignment.

**Figure 3.**
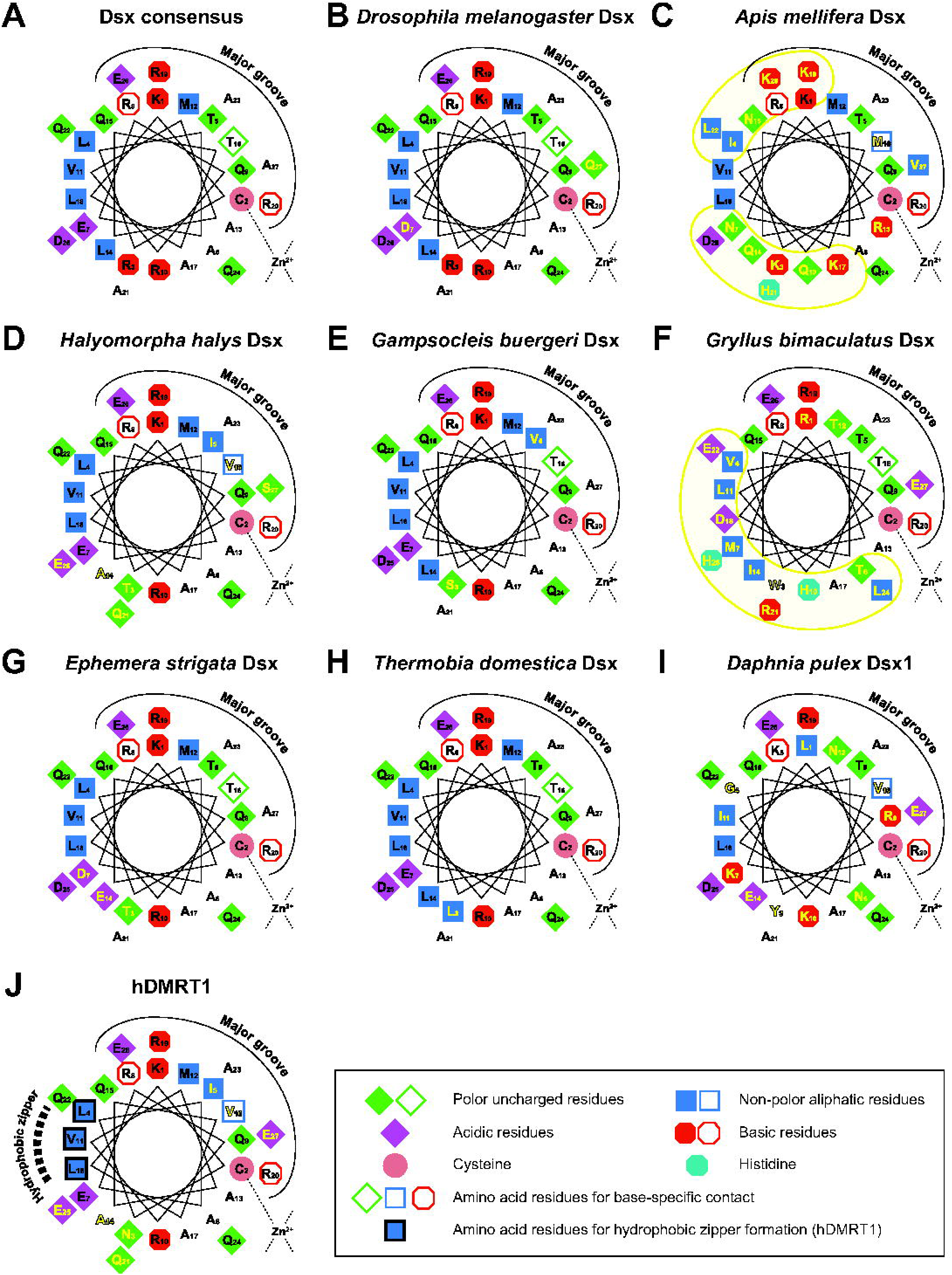
Helical wheel diagram representation of the major-groove recognition helices of various Dsx/DMRT proteins. (A) The consensus sequence of the major-groove recognition helices among insect and crustacean Dsx proteins (See Figure 2B). (B-I) The major-groove recognition helix of the Dsx homolog of various insect and crustacean species. (B) *Drosophila melanogaster*, (C) *Apis mellifera*, (D) *Halyomorpha halys*, (E) *Gampsocleis buergeri*, (F) *Gryllus bimaculatus*, (G) *Ephemera strigata*, (H) *Thermobia domestica*, and (I) *Daphnia pulex.* (J) The major-groove recognition helix of *Homo sapience* DMRT1 (hDMRT1). The properties of the amino acid residues are annotated with shapes and colors as described in the inset. Amino acid residues where substitutions occur compared to the consensus sequence are indicated by yellow letters. In (C) and (F), clusters of amino acid substitutions are highlighted with yellow shading. Amino acid residues required for base-specific contact are highlighted with outlined shapes with a white background. Amino acid residues involved in hydrophobic zipper formation are highlighted with black outlined squares with a blue background. Amino acid residues facing the major groove, coordinating zinc ions, required for base-specific contact, and involved in hydrophobic zipper formation in hDMRT1 are indicated based on the alignment shown in Figure 2B and Murphy et al. (2015).

To assess the degree of amino acid substitutions in the DM domain among insect Dsx homologs, a comprehensive multiple sequence comparison was performed, with a particular focus on the major-groove recognition helix (Figure 2B). The amino acid sequence of the major-groove recognition helix is generally conserved across insect species. However, significant amino acid substitutions were detected within two specific lineages: the suborder Apocrita (which includes wasps, bees, and ants; Hymenoptera) and the superfamily Grylloidea (‘true crickets’; Orthoptera: Ensifera). Among the seven Apocrita species included in this analysis, the major-groove recognition helix of the Dsx homolog in the honeybee *Apis mellifera* exhibited the highest substitution rate, with 51.9% of the sites (11 out of 21 residues) showing substitutions compared to the consensus sequence. Similarly, among the nine Grylloidea species included in this analysis, the major-groove recognition helix of the Dsx homolog in the tree cricket *Oecanthus longicauda* exhibited the highest substitution rate, with 60.7% of the sites (13 out of 21 residues) showing substitutions compared to the consensus sequence. In these two lineages, amino acid substitutions were not always shared between species, suggesting a high rate of substitution introduction into the major-groove recognition helix within the lineages. Following the two aforementioned lineages, moderate levels of amino acid substitutions (25.9-29.6% of the sites; 5-6 out of 21 residues) were observed in this region of Dsx homologs within Hemiptera. In Hemiptera, shared amino acid substitutions were observed among species, suggesting that a common ancestor of the lineage acquired these substitutions, which were conserved during the diversification of the lineage.

Regarding the *Gryllus* Dsx protein, it exhibited the second highest level of amino acid substitution compared to the consensus sequence (55.4% of the sites; 12 out of 21 residues) among all the Dsx homologs analyzed. The Cys-to-His substitution observed in the second zinc-binding site (site II) of *Gryllus* Dsx protein was conserved in all orthopteran Dsx homologs except for the mole cricket *Gryllotalpa orientalis* (Figure 2B). This suggests that the common ancestor of orthopteran insects acquired this substitution, which was later reverted in *Gryllotalpa*. Notably, amino acid residues crucial for dimer formation between DM domains—identified in the hDMRT1 protein by X-ray crystallography (Murphy et al., 2015)—were significantly substituted (10 out of 12 residues).

To assess the three-dimensional pattern of amino acid substitutions, the spatial positions of the amino acid residues within the major-groove recognition helix, as well as the interacting DNA and the zinc ion, are illustrated in a helical wheel diagram (Figure 3). The consensus sequence of the major-groove recognition helix among insect and crustacean Dsx homologs, along with the major-groove recognition helices of Dsx homologs from seven insect/crustacean species and that of *Homo sapiens* DMRT1 (hDMRT1), were depicted as helical wheel diagrams. In the helical wheel diagram, three amino acid residues that directly contact DNA bases are positioned at positions 8, 16, and 20. The amino acid residues at positions 8 and 20 are highly conserved Arg residues, whereas the residue at position 16 (Thr residue in the consensus) is substituted in some species, including *Apis mellifera*, *Halyomorpha halys*, and *Daphnia pulex*. In these species, the Thr residue is substituted with hydrophobic residues such as Met and Val. Amino acid substitution rates are low for amino acid residues facing the major-groove of interacting DNA except for the Dsx homologs in *Daphnia pulex*. In the Dsx homologs of *Apis mellifera* and *Gryllus bimaculatus*, which were selected as representative examples of two lineages with high substitution rates in the major-groove recognition helix, the substituted residues are clustered on the opposite side, away from the major groove (highlighted with yellow shading in Figure 3C and F), where a hydrophobic zipper is formed between the two DM domains of hDMRT1 proteins (Murphy et al., 2015). These data suggest that amino acid substitutions within the major-groove recognition helix can affect the DNA interaction and the dimer formation between DM domains, potentially altering the DNA-binding specificity of Dsx homologs.

### 3. Insect Dsx proteins share a conserved binding motif regardless of the substitutions in the major-groove recognition helix

To examine whether amino acid substitutions within the major-groove recognition helix alter the DNA-binding specificity of Dsx homologs, the DNA-binding specificity of seven DM domains representing insect phylogeny and sequence variation among Dsx homologs was determined using systematic evolution of ligands by exponential enrichment (SELEX). The DM domains of Dsx homologs (DsxDMs) of *Drosophila melanogaster, Ephemera strigata*, and *Thermobia domestica* were included in the analysis as representatives of Dsx homologs with major-groove recognition helices resembling the consensus sequence. The DsxDMs of *Apis mellifera*, *Gryllus bimaculatus*, and *Daphnia pulex* were included as representatives of Dsx homologs with highly substituted major-groove recognition helices. Additionally, a ‘back-mutated ‘*Gryllus bimaculatus* DsxDM, which contains a major-groove recognition helix from *Gampsocleis buergeri* that resembles the consensus sequence, was included in the analysis (*Gryllus bimaculatus* DsxDM^BM^) (Figure 4A). A random DNA Library containing a 26-bp random sequence was selected based on its affinity for each recombinant DsxDM protein. For each DsxDM protein, DNA libraries from eight rounds of SELEX were sequenced, and representative sequence motifs were identified from each library.

Figure 4B shows the representative sequence motifs for each DsxDM protein from SELEX rounds 3 and 4. To my surprise, the seven DsxDM proteins included in the analysis all recognize the same 7-nt palindromic motif, ACAyTGT, despite the substitutions in the recognition helix. In the cases of *Ephemera strigata* DsxDM and *Gryllus bimaculatus* DsxDM^BM^, the 7-nt motif further extends to a 9-nt motif: GCA<u>ACAyTGT</u> or <u>ACAyTGT</u>TGC (with the core ACAyTGT motif underlined). These results are consistent with those of previous studies conducted on *Drosophila melanogaster* Dsx protein, *Caenorhabditis elegans* Mab-3 protein, and vertebrate DMRT family proteins (Erdman et al., 1996; Yi & Zarkower, 1999; Murphy et al., 2007). Most notably, the results clearly demonstrate that naturally occurring substitutions found within the major-groove recognition helix hardly alter the DNA-binding specificity of Dsx homologs.

### 4. The *dsx* homolog in crickets is translocated from an autosome to the X chromosome

Later in this research, I generated knockout lines of the *Gryllus dsx* gene using CRISPR-Cas9 genome editing. During the establishment of the knockout line, I observed that the *Gryllus dsx* gene is linked to a sex chromosome. Since only scaffold-level genomic information is currently available for *Gryllus bimaculatus*, I investigated whether the *Gryllus dsx* gene is located on an autosome or the X chromosome by comparing the copy number of the *Gryllus dsx* gene in males and females via qPCR.

Three cricket species (*Gryllus bimaculatus*, *Teleogryllus commodus*, and *Acheta domestica*), as well as a katydid *Gampsocleis buergeri* and a migratory locust *Locusta migratoria*, were included in this analysis (Figure 5A). To ensure that the measurement of gene copy number was accurate, a *glucose-6-phosphate dehydrogenase* (*G6PD*) gene, which is known to be located on the X chromosome in mole cricket (Rao & Padmaja, 1992), was used as a positive control. Among the five orthopteran insects tested, the copy number of the *dsx* homolog in males was half that of females in all cricket species and a katydid, both of which belong to the suborder Ensifera. This finding indicates that the *dsx* genes in Ensifera species are located on the X chromosome. In contrast, in *Locusta migratoria* (suborder Caelifera), the copy number of the *dsx* homolog in males was the same as in females. This finding indicates that the *Locusta migratoria dsx* gene is located on an autosome, which is consistent with the location of the *dsx* homolog on chromosome 1 in *Schistocerca americana*, a grasshopper species for which chromosome-level genomic information is available (iqSchAmer2.1). To my knowledge, insect *dsx* homologs are typically located on autosomes, suggesting that the *dsx* gene was translocated from an autosome to the X chromosome in the common ancestor of the suborder Ensifera.

**Figure 5.**
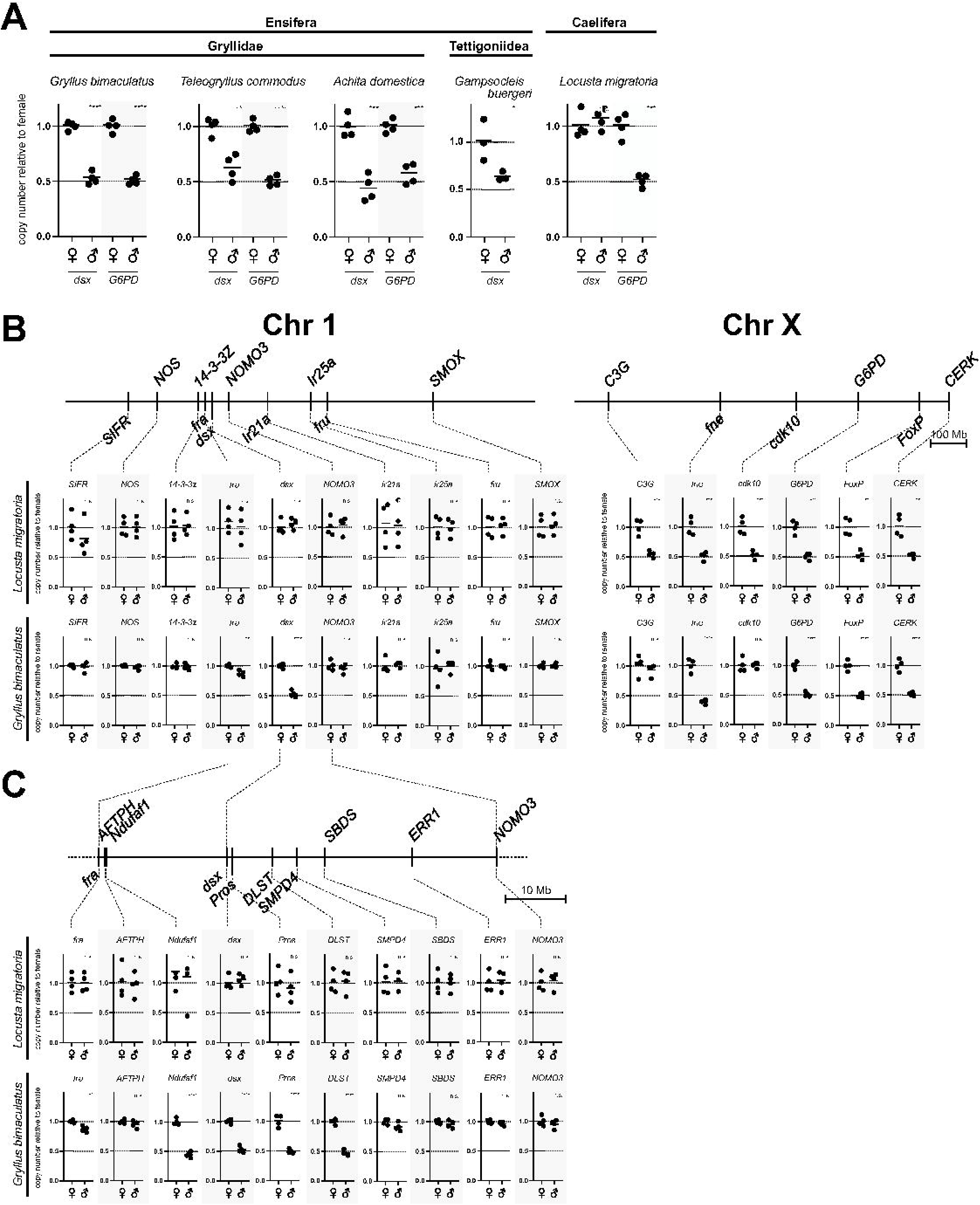
Gene copy number analysis to determine whether the *dsx* homolog and other GOIs are located on an autosome or the X chromosome. (A) Comparative gene copy number analysis of *dsx* homologs in five orthopteran species. The *G6PD* gene was included as a positive control for the X-linked gene. (B) Gene copy number analysis for a representative set of genes located on chromosome 1 and the X chromosome in the grasshopper *Schistocerca americana.* (C) Gene copy number analysis for genes located around the *dsx* gene in chromosome 1 of *Schistocerca american.* In (B) and (C), the gene copy number was measured in *Gryllus bimaculatus* and *Locusta migratoria*. The copy number of the GOIs was measured in males and females (n=4, respectively). The gene copy number for each individual was plotted, with the average value for females normalized to 1.0. Asterisks indicate statistical significance between the sexes (*, p<0.05; **, p<0.01; ***, p<0.001; ****, p<0.0001, n.s., p≥0.05).

Next, I investigated the type of chromosomal rearrangement underlying the translocation of the ancestral *dsx* gene from an autosome to the X chromosome in the Ensifera lineage. Without chromosome-level genomic information, chromosome synteny analysis is not applicable to this species. To investigate whether the translocation of the ancestral *dsx* gene was due to chromosome fusions or translocations of the genomic region surrounding *dsx*, I employed gene copy number analysis for a representative set of genes located on chromosome 1 and the X chromosome in the grasshopper *Schistocerca americana*, a ‘template ‘species of orthopteran chromosomes for this analysis. Quantitative PCR-based gene copy number analysis was conducted in the cricket *Gryllus bimaculatus* with the migratory locust *Locusta migratoria* serving as a control species (Figure 5B). In *Locusta migratoria*, the copy number of the 9 genes representing chromosome 1 of *Schistocerca americana* was the same between males and females, indicating that these genes are located on an autosome(s). The copy number of the six genes representing the X chromosome of *Schistocerca americana* was half in males compared to females, indicating that these genes are located on the X chromosome. In *Gryllus bimaculatus*, the copy number of eight out of the nine genes representing *Schistocerca americana* chromosome 1, excluding the *dsx* gene, was the same between males and females. The copy number of four out of the six genes representing the X chromosome in *Schistocerca americana*, excluding the *C3G* and *cdk10* genes, was half in males compared to females. The copy number of the *C3G* and *cdk10* genes was the same between males and females, indicating that these genes are located on an autosome(s) in *Gryllus bimaculatus*. These data suggest that the translocation of the ancestral *dsx* gene to the X chromosome was not due to chromosome fusion between the ancestral X chromosome and an autosome, as observed in some grasshopper lineages (Kawakami et al., 2011; Jetybayev et al., 2017). Instead, it appears to be a result of translocations of the genomic region surrounding the *dsx* gene.

Then, I estimated the size of the genomic region that was translocated to the X chromosome along with the *dsx* gene (Figure 5C). Again, the gene copy number analysis was conducted for seven genes located around the *dsx* gene in chromosome 1 of *Schistocerca american.* In *Gryllus bimaculatus*, the copy number of the three genes (*Ndufaf1*, *Pros*, and *DLST*) out of seven genes located around the *dsx* gene in *Schistocerca american* was half in males compared to females. In *Schistocerca americana*, the *Ndufaf1*, *dsx*, *Pros*, and *DLST* genes are tandemly aligned on chromosome 1, spanning a 27 kbp region. The *AFTPH* and *SMPD4* genes, located across this region, are 31 kbp apart in the *Schistocerca americana* genome. This suggests that the genomic region corresponding to this ∼30 kbp segment on chromosome 1 of *Schistocerca american* translocated from an autosome to the X chromosome in the common ancestor of the suborder Ensifera.

### 5. Knockout of the *Gryllus dsx* homolog resulted in morphological feminization

To investigate whether the *dsx* homolog is involved in sex determination in crickets, knockout lines of the *Gryllus dsx* gene were generated. CRISPR-Cas9 genome editing was employed to introduce indels into the coding exons common to all *Gryllus dsx* isoforms. After testing the efficiency of several gRNAs targeting the coding exons of *Gryllus dsx* gene, I selected a gRNA targeting exon 1 to induce indels specifically in the DM domain of the *Gryllus* Dsx protein. Cas9 RNPs targeting the *Gryllus dsx* gene were injected into the fertilized eggs of the Hokudai *gwhite* strain (G0 generation), which were then raised to adulthood. Some G0 adults exhibited a gynandromorph phenotype with abnormal forewing vein patterns (data not shown). These G0 adults were subsequently inbred. Female G1 crickets were raised to adulthood and backcrossed with the Hokudai *gwhite* strain, and were analyzed using T7E1 assay. As a result, nine G1 female crickets with heterozygous mutations, including indels in the DM domain of the *Gryllus* Dsx protein, were selected (Figure 6A). These G1 females were subsequently backcrossed to Hokudai *gwhite* strain to establish stable lines. G2 crickets were raised to adulthood and genotyped using a PCR-based method with primers designed to distinguish between the wild-type and mutant alleles. At this stage, I observed that I could collect female adults with only wild-type alleles, only mutant alleles, or both wild-type and mutant alleles. However, no male adults with mutant alleles were collected (data not shown). This could be due to either (1) male crickets harboring the mutated *Gryllus dsx* gene being developmentally lethal, or (2) the *Gryllus dsx* gene being X-linked. To test this, I conducted qPCR-based gene copy number measurements to determine if the *Gryllus dsx* gene is X-linked. As described above, the *Gryllus dsx* gene is indeed X-linked. This finding explains the unusual inheritance pattern of the mutated *Gryllus dsx* allele.

**Figure 6.**
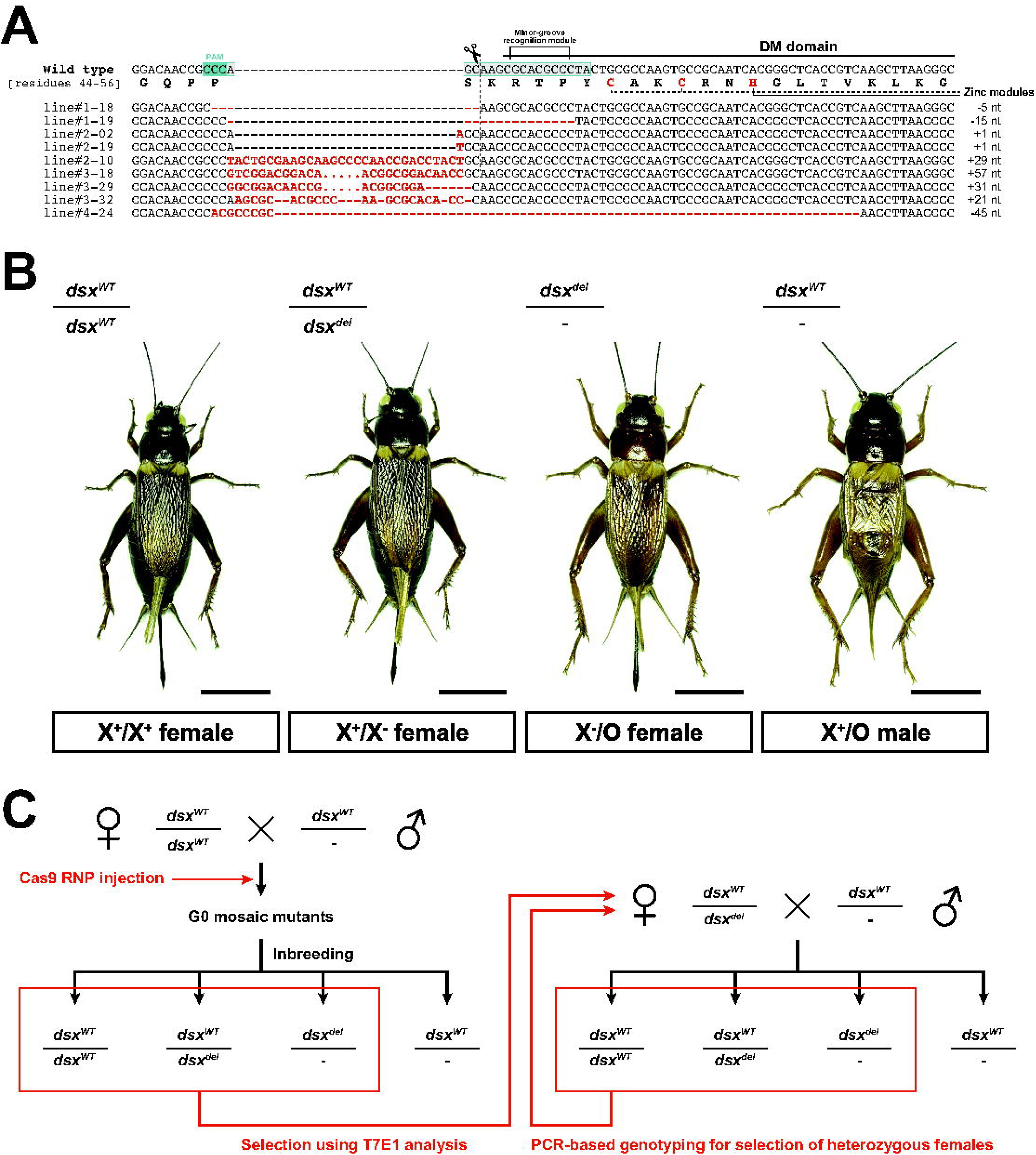
Generation of knockout lines of the *Gryllus dsx* gene using CRISPR-Cas9 system. (A) CRISPR-Cas9-induced indel mutations in the *Gryllus dsx* gene. A crRNA designed to cleave the 5 ‘region of the sequence encoding the DM domain was used. The *Gryllus dsx*^del^ mutant line was established using line #4-24, which harbors a 45-nt deletion in the coding sequence of the DM domain. (B) Phenotypes of *Gryllus dsx*^del^ mutants. One-week-old adults collected from the colony of the *Gryllus dsx*^del^ mutant line (line #4-24) were anesthetized with diethyl ether, photographed, and their genotypes were determined. Scale bars = 1 cm. Note that this species exhibits variations in body size. (C) Breeding strategy for *Gryllus dsx*^del^ mutant as an X-linked feminization allele. Crickets with female morphology were subjected to T7E1 assay at the G1 generation, followed by PCR-based genotyping in the subsequent generations. All breeding was conducted on the Hokudai *gwhite* mutant background.

Among nine *dsx* mutants, line #4-24, which has a 45-nt deletion, was selected for downstream analysis (*Gryllus dsx*^del^ mutant line). The mutated *Gryllus dsx* allele encodes a Dsx protein with a non-functional DM domain, as two Cys and one His residues required for zinc coordination are deleted. From a group of G2 adults, females with only mutant alleles or both wild-type and mutant alleles (heterozygous females) were collected as founders of subsequent generations. At this stage, I observed that females with only mutant alleles did not lay eggs.

Therefore, only heterozygous females were used to maintain the *Gryllus dsx*^del^ mutant line by backcrossing with Hokudai *gwhite* males (Figure 6C).

Figure 6B shows the morphology of adult crickets obtained from the colony of the *Gryllus dsx*^del^ mutant line (line #4-24), which was maintained by backcrossing heterozygous females with Hokudai *gwhite* males. The crickets are subdivided into four groups according to their genotype and morphology: (1) crickets with a pair of X chromosomes carrying the wild-type *Gryllus dsx* allele (*dsx^WT^*), exhibiting female morphology (*dsx^WT^*/*dsx^WT^*; X^+^/X^+^ females), (2) crickets with one X chromosome carrying the wild-type *Gryllus dsx* allele and another X chromosome carrying the mutated *Gryllus dsx* allele (*dsx^del^*), exhibiting female morphology (*dsx^WT^*/*dsx^del^*; X^+^/X^−^ females), (3) crickets with one X chromosome carrying the wild-type *Gryllus dsx* allele, exhibiting male morphology (*dsx^WT^*/-; X^+^/O males), and (4) crickets with one X chromosome carrying the mutated *Gryllus dsx* allele, exhibiting female morphology (*dsx^del^*/-; X^−^/O females). X^−^/O females have forewings with a female-type wing vein pattern and an ovipositor, morphologically identical to wild-type females and X^+^/X^−^ females (Figure 6C). This result indicates that the knockout of *Gryllus dsx* gene resulted in morphological feminization, supporting the idea that the *Gryllus dsx* gene functions as a masculinizing factor in crickets, which is consistent with the findings in the other hemimetabolous and ametabolous insects (Wexler et al., 2019; Chikami et al., 2022).

### 6. Knockout of the *Gryllus dsx* homolog did not alter sexual orientation in adult social behavior

As mentioned above, X^−^/O females were morphologically identical to wild-type females, but did not lay eggs when paired with adult males. During the breeding experiment, I observed that some X^−^/O females displayed abnormal social behaviors toward conspecifics, such as vibrating their forewings and aggressively chasing males—behaviors typically associated with male-specific social behavior in crickets. This observation raises a hypothesis: although the knockout of the *Gryllus dsx* gene resulted in morphological feminization, the sexual orientation in social behavior remains male-oriented. To test this hypothesis, I conducted a behavioral experiment comparing the social behavior of wild-type females (i.e., X^+^/X^+^ females), X^+^/X^−^ females, and X^−^/O females (Figure 7). A female cricket from the colony of the *Gryllus dsx*^del^ mutant line and a male or a female from the Hokudai WT strain, both 1 week old after adult emergence and isolated for three days, were subsequently introduced into an arena and allowed to interact for >5 minutes. Behavioral observations were conducted in a blind manner, without knowledge of the genotypes of the subjects. The behaviors of the female crickets were categorized into nine types: (1) Copulation, (2) Courtship, (3) Male-type aggression, (4) Female-type aggression, (5) Chasing, (6) Antennal contact, (7) Kicking, (8) Withdrawal, and (9) Being chased (criteria are summarized in Table 3; see Supplementary movies). After categorizing the behaviors, the female crickets were genotyped, and the behavioral data were divided into three groups based on genotype.

**Figure 7.**
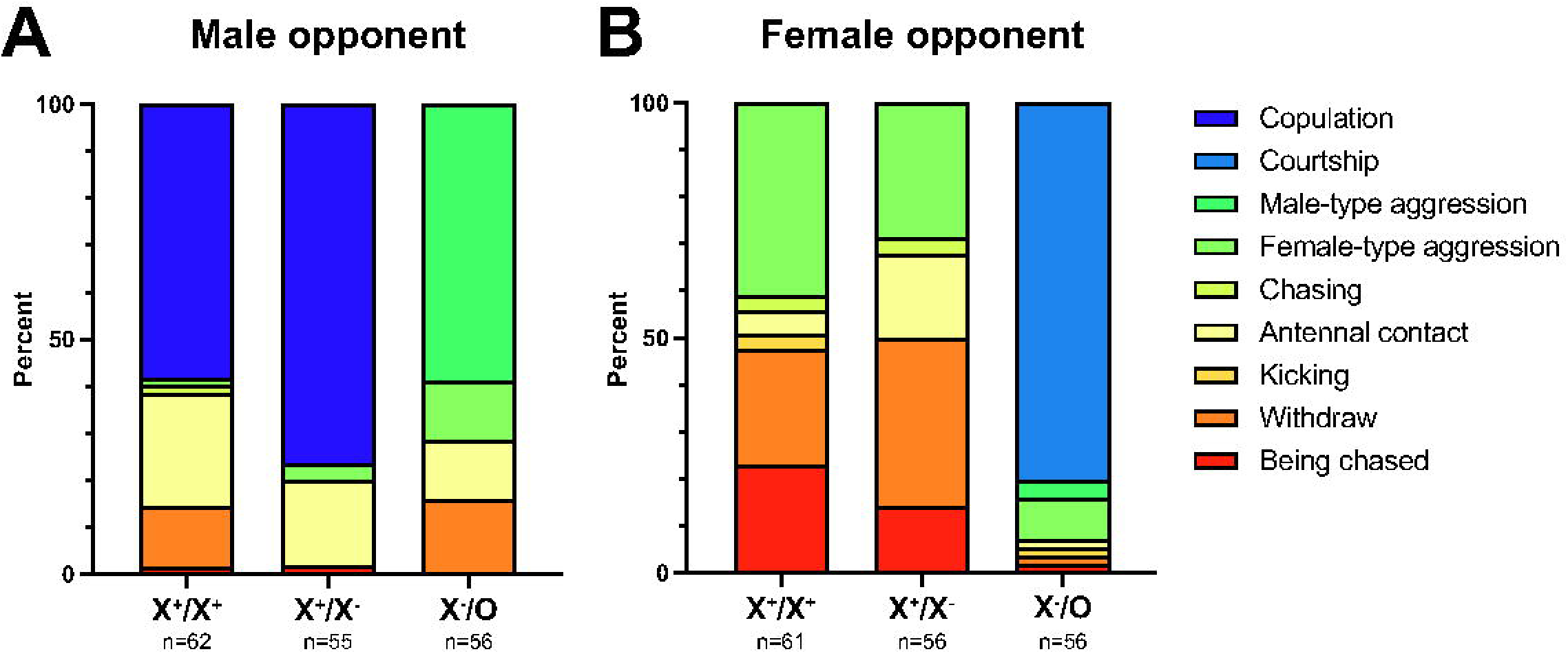
Social behavior of female adults of the *Gryllus dsx^del^* mutant. Behavioral responses of female adult crickets from the *Gryllus dsx^del^* mutant line when exposed to male or female adult wild-type crickets. The behavior of crickets was categorized into nine groups based on the criteria described in Table 3. (A) Behavioral responses to male adult crickets. (B) Behavioral responses to female adult crickets. The proportions of individuals exhibiting each behavioral category are represented in the stacked bar graphs.

Wild-type male adult crickets typically displayed courtship behavior towards X^+^/X^+^ females, X^+^/X^−^ females, and X^−^/O females. Eventually, 58% of X^+^/X^+^ females and 76% of X^+^/X^−^ females accepted the males for copulation. In contrast, 71.4% of X^−^/O females displayed agonistic behavior, with over 80% of these displaying male-type aggression (i.e., aggressive engagement with mandible spreading and/or wing raising), accounting for 58.9% of the total observations. None of the X^−^/O females copulated with wild-type males, and neither X^+^/X^+^ females nor X^+^/X^−^ females displayed male-type aggression to wild-type males (Figure 7A). When a wild-type female and a X^+^/X^+^ female or a X^+^/X^−^ female were introduced into the arena, they typically displayed territorial behavior, starting with weak agonistic interactions. This interaction could escalate to female-type aggression (i.e., aggressive engagement without mandible spreading and wing raising) or continuous chasing behavior. However, when a wild-type female and a X^−^/O female were introduced into the arena, the X^−^/O female predominantly displayed courtship behavior toward the wild-type female (80.4% of the total observations). The X^−^/O female rarely displayed agonistic behavior toward the wild-type female (12.5% of the total observations), and neither X^+^/X^+^ females nor X^+^/X^−^ females displayed courtship behavior toward the wild-type females (Figure 7B). These data strongly support the idea that the knockout of the *Gryllus dsx* gene resulted in morphological feminization without altering the sexual orientation of individuals, which remains determined by sex chromosome composition.

## Discussion

To understand the evolution of sex-determination mechanisms in the insect nervous system, a series of studies on the *dsx* gene was conducted in the two-spotted cricket, *Gryllus bimaculatus*. The *Gryllus dsx* gene had a different gene organization compared to previously reported *dsx* homologs and showed a unique sex-specific expression pattern. The *Gryllus* Dsx protein contained more amino acid substitutions in the DNA-binding DM domain than Dsx homologs in other insect lineages. Nevertheless, comparative structural and biochemical analyses revealed that the DM domain of the *Gryllus* Dsx proteins can perform DNA-binding function homologous to those in other insects. I also found that the *dsx* gene has been translocated to the sex chromosome in the cricket lineage. Furthermore, the knockout of the *Gryllus dsx* gene resulted in morphological feminization, indicating that *Gryllus dsx* is responsible for morphological masculinization.

Surprisingly, these feminized males retained male-type social behavior, suggesting that while the *Gryllus dsx* gene controls morphological masculinization, it does not play a central role in neuronal sex determination.

In the present study, I revealed that the cricket *dsx* gene has a unique genetic structure that is not found in other insect *dsx* homologs. In the fruit fly *Drosophila melanogaster*, the exon encoding the female-specific C-terminal tail of the Dsx^F^ isoform (exon 4) positions between the exon encoding the common UBA domain (exon 3) and the exons encoding the male-specific C-terminal of the Dsx^M^ isoform (exons 5 and 6). In the two-spotted cricket *Gryllus bimaculatus*, the exon encoding the C-terminal tail of the *Gryllus Dsx^F^* isoform (exon 7) is located furthest 3 ‘from the gene body. To my knowledge, the gene organization of most insect *dsx* homologs resembles that of *Drosophila* (Ohbayashi et al., 2001; Cho et al., 2007; Salvemini et al., 2011; Kijimoto et al., 2012; Shukla & Palli, 2012; Wexler et al., 2019; Baral et al., 2019), with female-specific exon(s) positioned between the common UBA exon(s) and the exon(s) encoding the male-type C-terminal tail. This indicates that the origin of the female-specific isoforms of *Gryllus dsx* differs from that of other insects.

In the brain of cricket, *Gryllus dsx* isoforms show sex-biased but not sex-restricted expression (Figure 1C). The molecular and cellular mechanisms underlying sex-biased splicing regulation remain unclear. One possibility is Tra/Tra2-dependent splicing regulation, similar to that in *Drosophila* (Hoshijima et al., 1991). In *Drosophila*, Tra2 binding motifs are located within exon 4, which encodes the female-specific C-terminal tail. In female cells, the Tra/Tra2 complex binds to the pre-mRNA of *Drosophila dsx*, leading to the inclusion of exon 4 during splicing. Conversely, in male cells, where functional Tra protein is not expressed, Tra fails to recruit the splicing machinery to the Tra2 binding motifs in exon 4, resulting in the exclusion of exon 4. This Tra/Tra2-dependent splicing regulation enables sex-restricted expression of different *dsx* isoforms in *Drosophila*.

Currently, the *Gryllus dsx* locus in the published genomic data (Gbim_1.0) contains many gaps in the intronic regions, making it difficult to determine whether Tra2 binding motifs are present in the *Gryllus dsx* gene. Further investigations are needed to understand the sex-biased alternative splicing mechanism of the *Gryllus dsx* gene.

Metazoan DMRT family proteins, except for Hexapoda Dsx homologs, lack both the UBA domain and the isoform-specific C-terminal tail (Price et al., 2015; Panara et al., 2019). This suggests that the UBA domain-dependent dimerization and transcription regulation via the isoform-specific C-terminal tail are unique features of insect Dsx homologs. The isoform-specific C-terminal tail of *Drosophila* Dsx^F^ contains a sequence motif for interaction with the cofactor Intersex (IX) (Garrett-Engele et al., 2002; Yang et al., 2008). This motif is conserved among Dsx^F^ isoforms in most holometabolous insects except for some hymenopterans, but not in polyneopteran insects, including the German cockroach *Blattella germanica* and *Gryllus bimaculatus* (Chikami et al., 2022), suggesting that the Dsx^F^ isoform in polyneopteran insects may not regulate transcription through interaction with the IX protein. This is consistent with the results of RNAi studies on *dsx* homologs in some ametabolous and hemimetabolous insects, where downregulation of *dsx* homologs led to male-to-female transformation, but not *vice versa* (Wexler et al., 2019; Chikami et al., 2022). Further biochemical and genetic studies are required to determine whether the ancestral Dsx^F^ is a non-functional isoform produced as a by-product of alternative splicing, or whether it has a transcriptional regulatory function independent of IX interaction.

Unlike the Dsx^F^ isoform in holometabolous insects, the molecular and cellular functions of the male-specific C-terminal tail of the Dsx^M^ isoform are poorly understood. The amino acid sequences of the male-specific C-terminal tail are moderately conserved across ametabolous to holometabolous insects (Figure 1B). The phylogenetic distribution of the Dsx^M^ isoform supports the idea that the ancestral Dsx acquired the male-type C-terminal tail at the early stages of insect evolution, and that this tail contributes to transcriptional regulation through a conserved, yet unknown mechanism across insect species. Further biochemical and genetic analyses are needed to elucidate the mechanisms of transcriptional regulation mediated by the male-specific C-terminal tail of insect Dsx^M^ protein.

In addition to the aforementioned Dsx^F^ and Dsx^M^ isoforms, the *Gryllus dsx* gene encodes an additional isoform, *Gryllus* Dsx^COM^, which is generated by intron retention of the UBA domain encoding exon (exon 4). In the firebrat *Thermobia domestica*, the female-specific isoform of *dsx* homolog is produced by intron retention of the UBA domain encoding exon (Chikami et al., 2022). In the *Gryllus dsx^del^* mutant, all three Dsx isoforms lose DNA-binding activity due to the deletion of zinc-coordinating residues in the DM domain. Therefore, the isoform-specific contributions to sex determination by the *Gryllus dsx* gene remain unclear. In *Drosophila*, the sex-specific *dsx* isoforms are exclusively expressed in males and females and are thus responsible for sex-specific gene expression regulation, respectively (Arbeitman et al., 2016). In the water flea *Daphnia magna*, the *dsx1* gene is restrictedly expressed in the male-specific structures and is involved in the development of male-specific traits (Kato et al., 2011). In crickets, I report for the presence of *dsx* isoforms that do not show sex-restricted expression. It is possible that the accelerated molecular evolution of the *dsx* gene in Ensifera, which will be discussed later, may have contributed to the complex isoform configuration in the *Gryllus dsx* gene. Further investigations are needed to elucidate the molecular and cellular mechanisms underlying the weak or non-sex-biased expression of *Gryllus dsx^M^* and *dsx^COM^* isoforms, as well as their functional significance and evolutionary origins.

In addition to the unique isoform configuration of the *Gryllus dsx* gene, it also appears to function as an X-linked masculinization factor, which was confirmed by gene copy number analysis and knockout study. Comparative gene copy number analysis across five orthopteran species revealed that the autosome-to-X chromosome translocation of the *dsx* gene in the common ancestor of Ensifera. In parallel with my efforts, Zhang et al. (2021) reported the presence of a *dsx* homolog on the X chromosome in the Hawaiian field cricket *Teleogryllus oceanicus*. Unfortunately, since chromosome-level genomic information for this species has not yet been published, I employed gene copy number analysis again in *Gryllus bimaculatus* to map out the genomic regions that have translocated along with the *dsx* gene. The result demonstrated that the genomic region corresponding to the ∼30 kbp segment on chromosome 1 of *Schistocerca americana*, where the *Ndufaf1*, *dsx*, *Pros*, and *DLST* genes are tandemly aligned, translocated from an autosome to the X chromosome in the common ancestor of the suborder Ensifera. To understand in detail how this autosome-to-X chromosome translocation of the *dsx* gene occurred, a comparative synteny analysis using chromosome-level genomic information from various orthopteran species is required.

As shown in Figure 2B, the major-groove recognition helix of insect Dsx homologs is highly conserved in almost all lineages of insects, except for the suborder Apocrita (Hymenoptera) and the superfamily Grylloidea (Orthoptera: Ensifera). Given that DMRT family proteins, including insect Dsx homologs, recognize a common binding motif and that the loss of Dsx function significantly disrupts sex determination, it is intriguing that the accumulation of amino acid substitutions in the major-groove recognition helix restrictedly occurred in particular insect lineages, such as Apocrita and Grylloidea. An interpretation from the point of view of evolutionary biology is that the autosome-to-X chromosome translocation of the *dsx* gene in Ensifera might enhance the rate of molecular evolution of the gene. As revealed in this study and previous research (Wexler et al., 2019; Chikami et al., 2022), *dsx* homologs in basal insects function as a masculinizing factor, indicating that this gene can be considered a typical male-beneficial gene.

Theoretical biology predicts that alleles beneficial to a particular sex are more likely to accumulate on sex chromosomes. Classical theory suggests that the X chromosome should favor female-beneficial alleles due to its sex-biased inheritance and greater time spent in females, while autosomes, which have an unbiased mode of inheritance, should equally favor alleles beneficial to both sexes (Rice, 1984; Mank, 2009; Mullon et al., 2012; Hitchcock & Gardner, 2020). Recent studies suggest that the X chromosome may more favor male-beneficial alleles than autosomes (Patten & Haig, 2009; Fry, 2010; Fry, 2013; Jaquiéry et al., 2013). Nevertheless, the autosome-to-X chromosome translocation of the *dsx* gene in the common ancestor of Ensifera drastically altered the genomic environment and inheritance dynamics of the chromosome where the *dsx* gene is located, likely accelerating the molecular evolution of the *dsx* gene in Ensifera, especially in Grylloidea. Similarly, in Hymenoptera, where sex is determined by haplodiploidy, the genome is transmitted in a pattern similar to that of the X chromosome in the XX:XO system. Therefore, the accumulation of amino acid substitutions in the major-groove recognition helix in Apocrita can be explained by similar evolutionary forces.

Another unique feature found in *Gryllus* and other orthopteran Dsx homologs is the amino acid substitution in the zinc module of the DM domain. Orthopteran Dsx homologs, except for *Gryllotalpa orientalis* Dsx, contain non-canonical CCHC-type and H<u>H</u>CC-type zinc-binding sites (with the position of the Cys-to-His substitution underlined). Both Cys and His residues are found in zinc finger proteins as zinc-coordinating residues, but His has a bulkier imidazole side chain compared to Cys and is less prone to oxidation. Zhang et al. (2006) conducted a mutational study on the DM domain of the *Drosophila* Dsx protein and reported that a Cys-to-His substitution at Cys68 (the residue corresponding to the Cys where the substitution occurs in orthopteran Dsx homologs) resulted in a loss of function. Despite this previous report, the Cys-to-His substitution may not disrupt the proper conformation of the DM domain in orthopteran Dsx homologs. This is supported by the successful use of SLELX with DM domains containing either canonical or Orthoptera-specific non-canonical zinc-binding sites. The Cys-to-His substitution requires a two-step point mutation in the first and second positions of the Cys encoding codon (TGT/TGC to CAT/CAC). At the first step of these point mutations, the Cys residue is substituted with Tyr or Arg, which do not coordinate with a zinc ion. This indicates that the intermediate Dsx protein loses the correct conformation of the DM domain. Despite a recent report on Malacostraca Dsx homologs, where a Cys-to-His substitution occurs at the same position as in orthopteran Dsx homologs (Zhang et al., 2023), most of the ∼3,000 DM domain-containing proteins registered in the Pfam database have intertwined CCHC-type and HCCC-type zinc-binding sites. This suggests that Cys-to-His substitutions in zinc-coordinating residues are rare in metazoan DMRT family proteins. The phylogenetic distribution of Cys-to-His substitutions in the DM domain of orthopteran Dsx homologs supports the hypothesis that this amino acid substitution originated in the common ancestor of orthopterans and subsequently spread through its descendant lineage. In the common ancestor of orthopterans, the ancestral *dsx* gene would have been located on an autosome, and the presence of a spare gene would have allowed the Cys-to-His substitution via two-step point mutations. However, the Dsx homolog in *Gryllotalpa orientalis* possesses a DM domain with canonical intertwined CCHC-type and HCCC-type zinc-binding sites, suggesting that the substituted His residue was back-mutated to Cys after the autosome-to-X chromosome translocation of the *dsx* gene in this lineage. This may be another evidence of an accelerated molecular evolution of the *dsx* gene in Ensifera.

The most significant finding of this study is that the *Gryllus dsx* gene functions as a masculinizing factor for morphological traits, but it does not determine sexual orientation in social behavior. Interestingly, the knockout of the *Gryllus dsx* gene did not affect sex-specific behavioral patterns. When X^−^/O females engaged with male adults, they spread their mandibles in the same way as wild-type males. X^−^/O females also displayed stridulation associated with male-specific behaviors, such as courtship. This suggests that not only the neural circuits that determine male-type sexual orientation, but also those that generate the behavioral patterns and sequences of behavioral elements that make up male sexual behavior, are retained functional in X^−^/O females.

This finding may explain the morphological and behavioral characteristics of ‘flatwing’ crickets (*Teleogryllus oceanicus*) in the Hawaiian Islands. The introduction of the parasitoid fly *Ormia ochracea* has led to the evolution of flatwing crickets, which lack sound-producing structures on their forewings. Since parasitic flies are attracted to the songs of adult male crickets, the silent morph is advantageous in avoiding parasitic flies. Although male flatwing crickets do not produce sound, they exhibit wing movement patterns similar to those of normal males (Schneider et al., 2018), suggesting that they have not, at least for the time being, lost the behavior characteristic of adult male crickets. The first attempt to identify the mutation responsible for the flatwing phenotype indicated that flatwing phenotype is caused by an X-linked mutation (Pascoal et al., 2014). Further research has narrowed down the responsible genomic region and revealed that a candidate mutation resides in a chromosomal region containing the *dsx* and *Pros* genes (Zhang et al., 2021; Rayner et al., 2024). This region corresponds exactly to the region where the autosome-to-X chromosome translocation had occurred in the common ancestor of Ensifera. It was also reported that the expression level of the *dsx* homolog in the wings of male flatwing crickets is reduced (Zhang et al., 2021; Rayner et al., 2024). Although the *dsx* homolog has not yet been identified as the causative gene for the flatwing phenotype, the results of the present study suggest that mutations in the regulatory regions of *dsx* might be present in flatwing crickets. As reported in this study, a loss-of-function mutation of the *dsx* homolog in crickets would result in complete morphological feminization, whereas mutations in the transcriptional regulatory elements responsible for regulating expression in the male wings may result in wing-restricted feminization. The present study also suggests that even if the function of the *dsx* gene is disrupted in the nervous system of flatwing crickets, it is unlikely to affect male social behavior. In addition, flatwing-like phenotypes are reported to have independently evolved multiple times in the Hawaiian Islands (Pascoal et al., 2014; Pascoal et al., 2016). Although there is no evidence that these are all due to mutations in the same causative gene (i.e., *dsx* homolog), it is not difficult to imagine that the accelerated molecular evolution of the cricket *dsx* gene shown in this study would result in a higher frequency of polymorphism in the *dsx* locus (and/or its surrounding genomic region) in the population. This would explain the independent origin of flatwing-like phenotype in different populations after the introduction of the parasitoid fly.

In *Drosophila*, extensive neurogenetic studies have revealed that neurons expressing the two sex-determination transcription factor genes, *dsx* and *fruitless*, regulate sexual orientation and the behavioral elements associated with sex-specific social behavior (Manoli et al., 2005; Demir & Dickson, 2005; Vrontou et al., 2006; Chan & Kravitz, 2007; Rideout et al., 2007; Kimura et al., 2008; Rideout et al., 2010; Dauwalder, 2011; Shirangi et al., 2016; Ishii et al., 2020; Han et al., 2022; Sato & Yamamoto, 2023). In some holometabolous insects, such as the fruit fly *Drosophila melanogaster*, the honeybee *Apis mellifera*, and the parasitic wasp *Nasonia vitripennis*, the *fruitless* transcript expressed from a promoter positioned furthest from the gene body (P1 promoter) undergoes sex-specific splicing (Ryner et al., 1996; Bertossa et al., 2009). I previously investigated whether the *fruitless* homolog is involved in neuronal sex determination in crickets and other hemimetabolous insects and found that the *fruitless* homolog in crickets lacks the P1-like promoter and does not undergo sex-specific splicing to produce sex-specific protein isoforms (Watanabe et al., 2019). This gene organization is likely conserved among other hemimetabolous insects; therefore, I proposed an evolutionary model of the insect *fruitless* homolog, as shown in Figure 8A. Together with the results of the present study, it is suggested that none of the known terminal sex-determination factors in insects contribute to the neuronal sex determination in crickets (Figure 8B, C). It is important to investigate whether the sex-specific splicing factor Tra, which regulates the sex-specific splicing of *doublesex* genes in many insects (Verhulst et al., 2010), contributes to sex determination in the cricket brain. RNAi-mediated down-regulation of the *tra* homolog in the female German cockroach *Blattella germanica* resulted in a complete switch of *dsx* splicing patterns from female to male, and the masculinization of female sexual behavior (Wexler et al., 2019). I have already confirmed that the *Gryllus tra* homolog expressed in the brain undergoes sex-specific splicing (data not shown). If Tra also regulates the orientation and pattern of sexual behavior in crickets, there must be an unknown neuronal sex determination factor downstream of Tra that undergoes sex-specific splicing in the cricket brain. Extensive studies are needed to elucidate the molecular and cellular mechanisms of neuronal sex determination in crickets.

**Figure 8.**
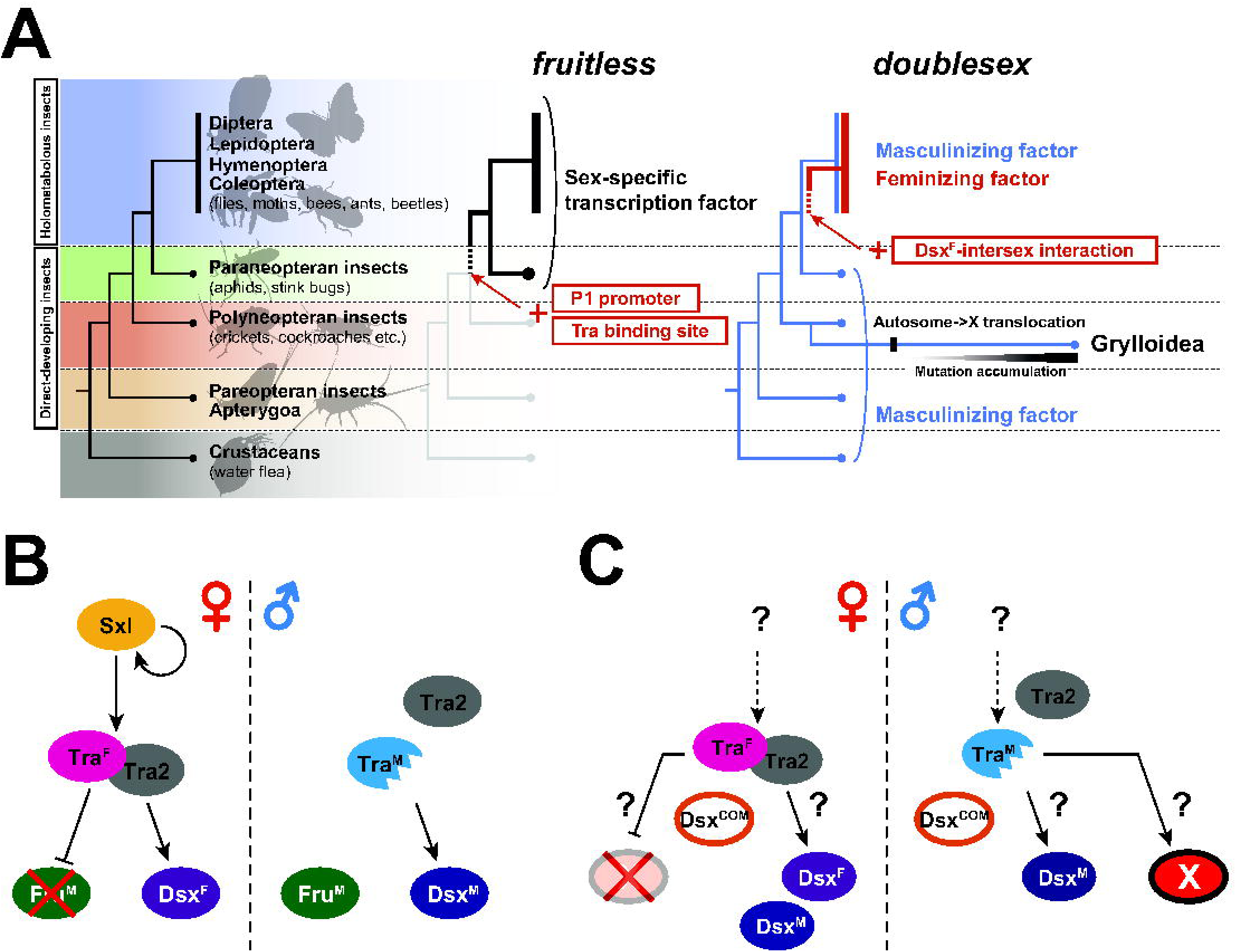
Evolution of neuronal sex determination system in insects. (A) The evolutionary hypothesis of two terminal differentiator genes in *Drosophila* neuronal sex determination cascade, *fruitless* and *doublesex*, across the evolution of insects. (B, C) Sex-determination cascade in the nervous system of *Drosophila melanogaster* (B) and *Gryllus* bimaculatus (C). (B) *Drosophila* neuronal sex determination relies on Tra/Tra2-mediated sex-specific splicing, and sexually dimorphic gene products of *dsx* and *fruitless* (Fru^M^). (C) Neither *doublesex* nor *fruitless* regulates neuronal sex determination in the nervous system of crickets. Instead, an unknown factor, X, plays a central role in neuronal sex determination in crickets. The role of Tra/Tra2-mediated sex-specific splicing in the neuronal sex determination has not yet been clarified.

## Author contributions

T.W. designed the study, performed experiments, analyzed the data, and wrote the manuscript.

## Acknowledgment

This research was supported by the Japan Society for the Promotion of Science, Grant Numbers JP19K06754 and JP23K05255 to T.W., and Akiyama Life Science Foundation research grant 2014 to T.W.

## Competing financial interests

The author declares that there are no conflicts of interest.

## Data availability

The high throughput sequencing data generated during the current study are available in BioProject (BioProject ID: PRJDB18769).

## Figure legends

**Supplementary Movie 1. Interaction between an X^+^/ X^+^ female (Hokudai *gwhite* background) and a wild-type male**.

**Supplementary Movie 2. Interaction between an X^−^/O female (Hokudai *gwhite* background) and a wild-type male**.

**Supplementary Movie 3. Interaction between an X^+^/ X^+^ female (Hokudai *gwhite* background) and a wild-type female**.

**Supplementary Movie 4. Interaction between an X^−^/O female (Hokudai *gwhite* background) and a wild-type female**.

**Supplementary Movie 5. Interaction between two X^−^/O females (Hokudai *gwhite* background)**

## Notes

### Competing Interest Statement

The authors have declared no competing interest.

